# *In vivo* KRAS G12D/V degradation mediated by CANDDY using a modified proteasome inhibitor

**DOI:** 10.1101/2021.04.23.441075

**Authors:** Satoshi Imanishi, Lijuan Huang, Shoko Itakura, Masamichi Ishizaka, Yoichi Iwasaki, Tomohiro Yamaguchi, Etsuko Miyamoto-Sato

## Abstract

“Undruggable” proteins, such as RAS proteins, remain problematic despite efforts to discover inhibitors against them. KRAS mutants are prevalent in human cancers. Recently, the KRAS G12C inhibitor have been clinically approved, but inhibitors for KRAS G12D/V are still under development. Here, we described the development of a novel chemical knockdown strategy, termed CANDDY (Chemical knockdown with Affinity aNd Degradation DYnamics). This strategy involves a CANDDY tag modified from a proteasome inhibitor inducing direct proteasomal degradation. We constructed TUS-007 as a multispecific small molecule tethered from a KRAS interactor and CANDDY tag to target KRAS G12D/V. We confirmed that the degradation by TUS-007 was independent of target ubiquitination. This allows to solve a laborious design of matchmaker in the current ubiquitination-dependent proteolysis technology. And TUS-007 successfully suppressed tumors due to in vivo degradation of KRAS G12D/V. The CANDDY technology could represent a simple and rational strategy to degrade currently “undruggable” proteins.

## Introduction

The majority (75%) of disease-causing proteins are “undruggable” (i.e., difficult to inhibit with small molecules). This difficulty reflects in the presence of smooth surfaces and lack of deep pockets, including proteins associated with cancer drivers and many interfaces for protein-protein interaction (PPI) (1, 2, 3). RAS family members (e.g., KRAS, HRAS, and NRAS) are most challenging to inhibit by small molecules (1, 4, 5, 6, 7). Mutations of KRAS, especially G12C, G12D, and G12V, are frequent in human cancers (7). The RAS protein activities depend on nucleotide loading in their GTP-binding pockets. The inhibition of this pocket has been attempted for nearly 40 years. However, progress has been hindered by the exceptionally high affinity between GTP and RAS proteins (1, 4, 8). While the inhibitor targeting KRAS G12C have been clinically approved recently (6, 9), inhibitors for KRAS G12D/V, which addresses greater clinical need, are still remained under development. These are important targets found in 95% of pancreatic cancers and 64% of colon cancers (7, 8). Inhibitors of the PPI between RAS and Son of Sevenless 1 (SOS1) (RAS-SOS inhibitor) have been investigated. Inhibitors that directly bind to RAS are not effective owing to their low affinity (5). Thus, the current inhibition technologies are not enough to be effective for undruggable proteins.

Pharmaceutical research on undruggable proteins, such as RAS, focuses on novel modalities instead of inhibition (10, 11, 12). Proteolysis technologies, using matchmakers between the target and E3 ligase or other ubiquitination related proteins such as molecular chaperons for inducing target degradations, are expected to be effective against undruggable proteins (13, 14, 15, 16, 17). However, matchmaker design has been hampered by the dependency on target ubiquitination (16). It is difficult to select a suitable E3 ligase for a target of interest because of limited knowledge about the mechanism underlying substrate recognition by E3 ligases. Even if the target has an established corresponding E3 ligase, there may be no available ligand for the E3 ligase. Moreover, target ubiquitination may not always induce proteolysis, as in the case of RAS, in which ubiquitination also regulates protein localization and activation (18). Despite such difficulties, an effective inhibitor (9) was used recently for a proteolysis inducer of KRAS G12C (19). However, no proteolysis inducer for KRAS G12D/V has been reported. Current proteolysis technologies that depend on ubiquitination have limited efficacy for undruggable targets (14). Therefore, to modulate diverse undruggable targets, the difficulties in matchmaking should be eliminated (13).

Here, we report successful KRAS G12D/V degradation, mediated by a novel tool named CANDDY (<u>C</u>hemical knockdown with <u>A</u>ffinity a<u>N</u>d <u>D</u>egradation <u>DY</u>namics). This innovative approach induces direct proteasomal degradation (i.e., chemical knockdown) of the target using a CANDDY tag derived from a proteasome inhibitor that lacks the site for inhibitory activity (inhibitor site). By CANDDY technology, chemical knockdown occurs without ubiquitination, so it can bypass the difficulties of designing a matchmaker in current proteolysis techniques (13). Using an inefficient target inhibitor, RAS/SOS inhibitor (5), we designed and synthesized a CANDDY molecule called TUS-007 for KRAS G12D/V. We demonstrated KRAS G12D/V chemical knockdown mediated by TUS-007 in cell-free, *in vitro*, and *in vivo* assays, and *in vivo* tumor suppression.

## Results

### A CANDDY molecule for targeting KRAS G12D/V proteins

CANDDY molecules are bispecific molecules constructed from two modules; a target interactor, which enables specific binding to the target, and a CANDDY tag, which induces proteasomal degradation (Fig. 1a). The CANDDY tag is essential in CANDDY technology and is a derivative lacking the inhibitor site of MLN2238 (MLN) (20), a clinical proteasome inhibitor (Fig. 1b and Text S1.). To design TUS-007, we employed a RAS-SOS inhibitor (5) (shown in Fig. 1c) which directly binds to KRAS G12D, KRAS G12V, and wild-types of KRAS and HRAS as the target interactor module. Since the binding to wild-type NRAS have not been reported, it was expected to avoid the severe toxicity observed in a pan-RAS inhibitor (21). RAS-SOS inhibitor was conjugated to the CANDDY tag using a NH_2_ linker (Fig. 1c and Text S1.).

**Fig. 1.**
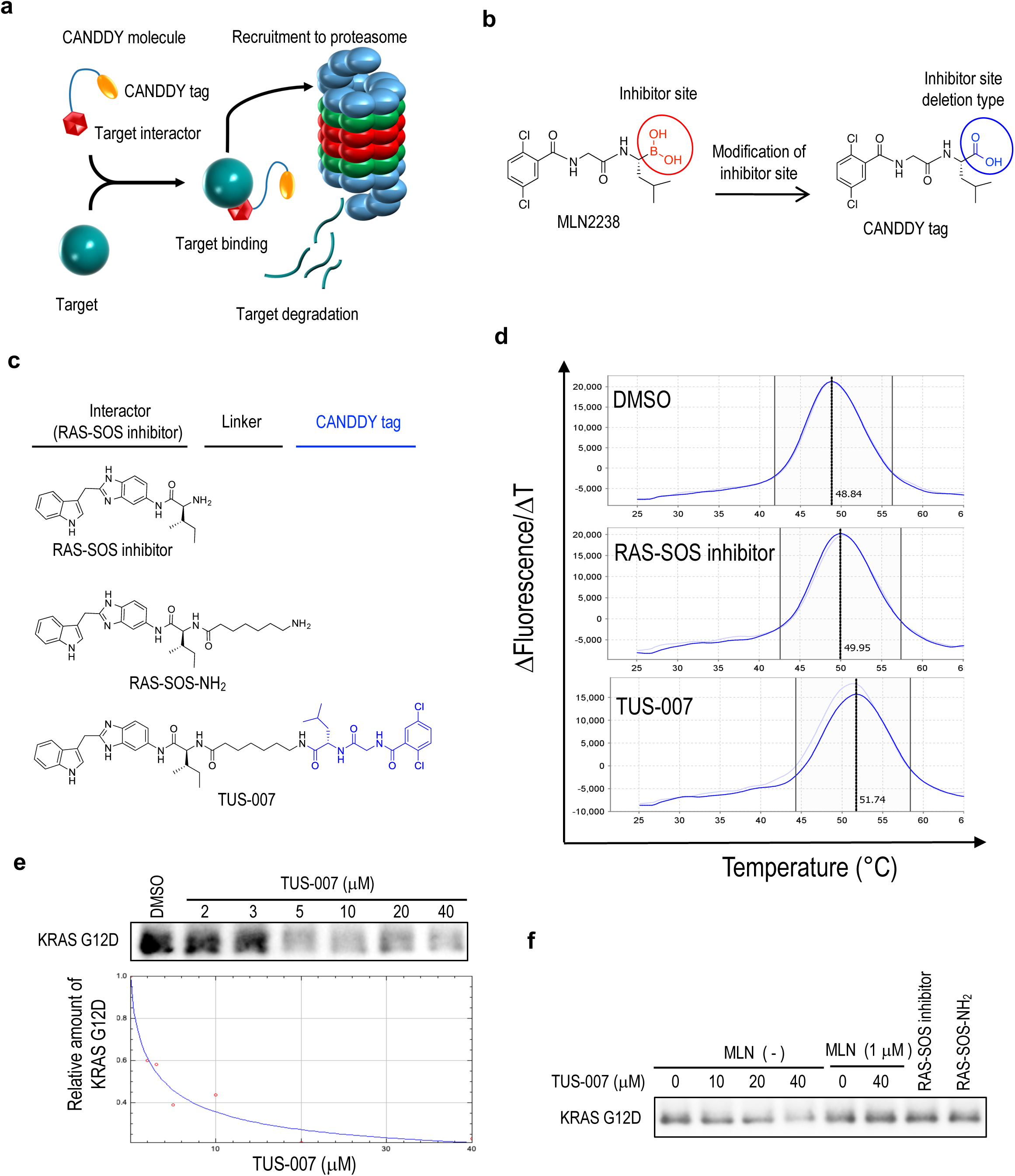
RAS chemical knockdown in cell free assays. **a,** Principle of the CANDDY technology. Each CANDDY molecule includes a target interactor (burgundy hexagon) and a CANDDY tag (yellow oval). The target protein is bound by the CANDDY molecule *via* the target interactor and is directly degraded by proteasome via the CANDDY tag. **b,** The CANDDY tag was synthesized by chemically inactivating the inhibitor site of MLM2238 (MLN). **c,** Structures of the RAS-SOS inhibitor (ref. 5), RAS-SOS-NH_2_, and TUS-007. **d,** KRAS G12D was mixed with DMSO, RAS-SOS inhibitor (4 μM) or TUS-007 (4 μM) and incubated under heating from 25 °C to 99 °C. The denature of KRAS G12D was monitored by the fluorescence. The typical curve of each group was shown. **e,** The correlation between relative amount of KRAS G12D and concentration of TUS-007 were approximated with Rodbard using the means of 3 independent examinations. The DC50 was estimated about 4 μM. **f,** TUS-007 induced KRAS G12D chemical knockdown by 26S proteasome in a cell free assay. KRAS G12D was incubated with 26S proteasome and agents as indicated for 3 h.

To compare the target binding of TUS-007 with that of the RAS-SOS inhibitor, we performed a thermal shift assay. Unexpectedly, KRAS G12D/V proteins incubated with TUS-007 were more resistant against heat treatment than those incubated with RAS-SOS inhibitor (Fig. S1a). Alternatively, in a fluorescence-based thermal shift assay, the T_m_ value of KRAS G12D incubated with TUS-007 was higher than that of KRAS G12D incubated with RAS-SOS inhibitor (Fig. 1d and Fig. S1b). These results suggest the higher affinity of TUS-007 to KRAS compared to RAS-SOS inhibitor. In addition, we confirmed that TUS-007 had no inhibitory activity against the catalytic β-subunits of proteasome (Fig. S2). This demonstrated that the CANDDY tag, modified from a proteasome inhibitor, hardly inhibit proteasome activity.

### Chemical knockdown of KRAS G12D/V free from ubiquitination by TUS-007

We attempted a chemical knockdown of KRAS G12D in the presence of 26S proteasomes and in the absence of any ubiquitination related proteins in a cell-free assay. Successful degradation was performed with 50% degradation concentration (DC50) at 4 μM (Fig. 1e). The chemical knockdown mediated by TUS-007 was counteracted by the presence of MLN (Fig. 1f). Additionally, RAS-SOS-NH_2_ (a synthetic intermediate without CANDDY tag, Fig. 1c) as a degradation-incompetent control failed to induce the chemical knockdown (Fig. 1f), demonstrating CANDDY tag is essential to induce the chemical knockdown. Although RAS-SOS inhibitor showed 80% inhibition of RAS-SOS PPI at 1 mM (5), DC80 of TUS-007 for KRAS G12D was estimated as 16 μM (Fig. 1e). It implied that the conjugation with CANDDY tag drastically improved the usefulness of RAS-SOS inhibitor. We also confirmed chemical knockdown of KRAS G12V by TUS-007 in a cell-free assay in the absence of any ubiquitination related proteins (Fig. S3). Furthermore, to obtain additional evidence of degradation independent of ubiquitination, we co-treated HeLa cells expressing GFP-KRAS G12D and DsRed with TUS-007 and MLN, E1 inhibitor (NAEi) or HSP70 inhibitor (VER-155008) (Fig. 2a). MLN counteracted the lowering of GFP signal induced by TUS-007 treatment, but NAEi or VER-155008 did not. These results demonstrated that chemical knockdown induced by TUS-007 was depend on proteasome and independent of ubiquitination processes.

**Fig. 2.**
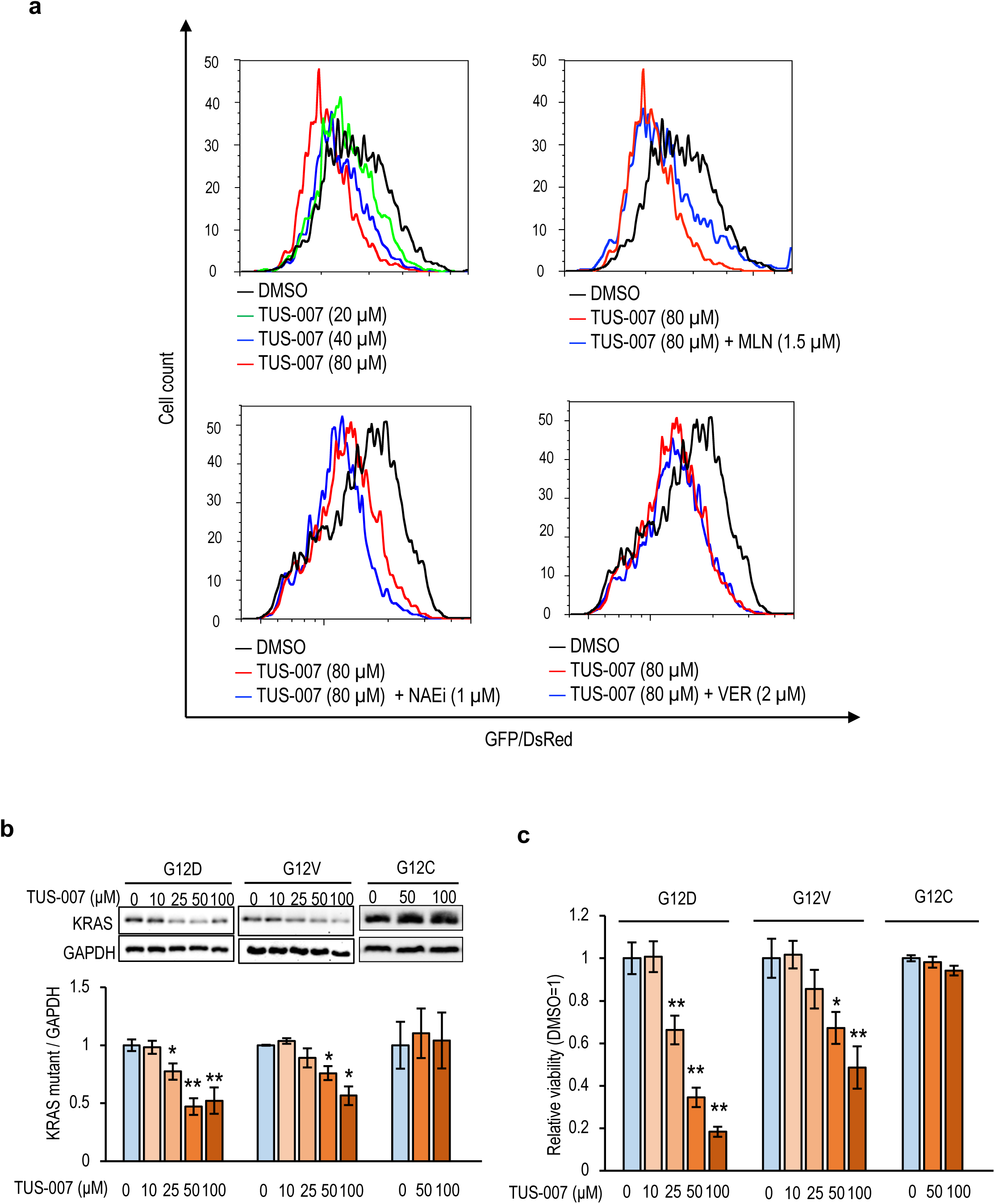
RAS chemical knockdown in cells. **a,** TUS-007 induced chemical knockdown was independent of ubiquitination. GFP fused KRAS G12D and DsRed were transiently expressed in HeLa cells. The cells were treated with TUS-007 in the presence of a proteasome inhibitor (MLN), a ubiquitination inhibitor (NAEi) and a molecular chaperon inhibitor (HSP70 inhibitor: VER-155008) for 24 h. **b & c,** RAS-less MEFs expressing KRAS G12D, G12V or G12C were incubated with TUS-007 for 72 h. Representative blots of KRAS are shown (**b**). The relative amounts of KRAS mutants normalized to GAPDH are shown in the bar graph (mean ± SEM; n = 3–4). The cell viability was also measured (mean ± SEM; n = 3–4) (**g**). *P < 0.05 and ** P < 0.01 vs. DMSO. TUS-007 induced chemical knockdown did not cause non-specific cytotoxicity.

### TUS-007 induced targets-selective chemical knockdown *in vitro*

Using an *in vitro* assay, we then evaluated the chemical knockdown and investigated the selectivity of TUS-007. We utilized RAS-less mouse embryonic fibroblasts (MEFs) expressing human KRAS G12D, G12V, or G12C. These MEFs do not proliferate in the absence of RAS (22). Immunoblotting analysis revealed that TUS-007 induced the chemical knockdown of KRAS G12D/V, but not G12C, in RAS-less MEFs (Fig. 2b). TUS-007 reduced the viability of RAS-less MEFs expressing KRAS G12D/V but did not affect the viability of the cells expressing KRAS G12C (Fig. 2c), confirming the selectivity of TUS-007 for KRAS G12D/V. We also assessed the selectivity of TUS-007 for the human RAS family, including wild-type KRAS, HRAS, and NRAS. TUS-007 did not affect the NRAS protein levels or the viability of RAS-less MEFs expressing NRAS but affected KRAS or HRAS levels and the viability of RAS-less MEFs expressing KRAS or HRAS (Fig. S4a and b). This result is consistent with the previously reported data on the selectivity of the RAS-SOS inhibitor (5). Therefore, it is expected not to induce intolerable toxicity previously observed with a pan-RAS inhibitor *in vivo* (21) since TUS-007 is not a pan-RAS degrader. Furthermore, the viability of RAS-less MEFs expressing KRAS G12C maintained even at 100 µM TUS-007 (Fig. 2b and c), indicating that TUS-007 might have no remarkable off-target toxicity in cell even at high concentrations. These results demonstrated the correlation between the degradation of targets and the viability of cells by the concentrations of TUS-007 without non-specific cytotoxic effect (Fig. 2b and c).

### The effects of TUS-007 on KRAS G12V-driven colon cancer

Approximately 64% of human colon cancers reportedly express KRAS mutants. KRAS G12V is found in nearly half of the patients and is correlated with poor prognosis (8). Therefore, we investigated whether TUS-007 can suppress the growth of human KRAS G12V-driven colon cancer cells. We conducted an experiment with the SW620-Luc KRAS G12V homozygous human colon cancer cell line (23). TUS-007 induced chemical knockdown of KRAS (Fig. 3a), accompanied by an increase in the annexin V-positive fraction in SW620-Luc cells (Fig. 3b and Fig. S5). However, RAS-SOS-NH_2_ and cetuximab did not induce apoptosis (Fig. 3b and Fig. S5). In contrast, treatment with TUS-007 did not result in significant changes in the annexin V-positive fraction in HT29-Luc RAS-independent colon cancer cells (Fig. 3c and Fig. S6). Alternatively, the apoptosis induction in SW620-Luc cells was confirmed by caspase 3/7 activation (Fig. 3d). Importantly, TUS-007 induced the apoptosis in SW620-Luc cells at the same concentration at which the chemical knockdown of KRAS was significant (Fig. 3a, b and Fig. S7). These findings indicated that TUS-007 selectively induces apoptosis in SW620-Luc cells by chemical knockdown of KRAS G12V *in vitro*.

**Fig. 3.**
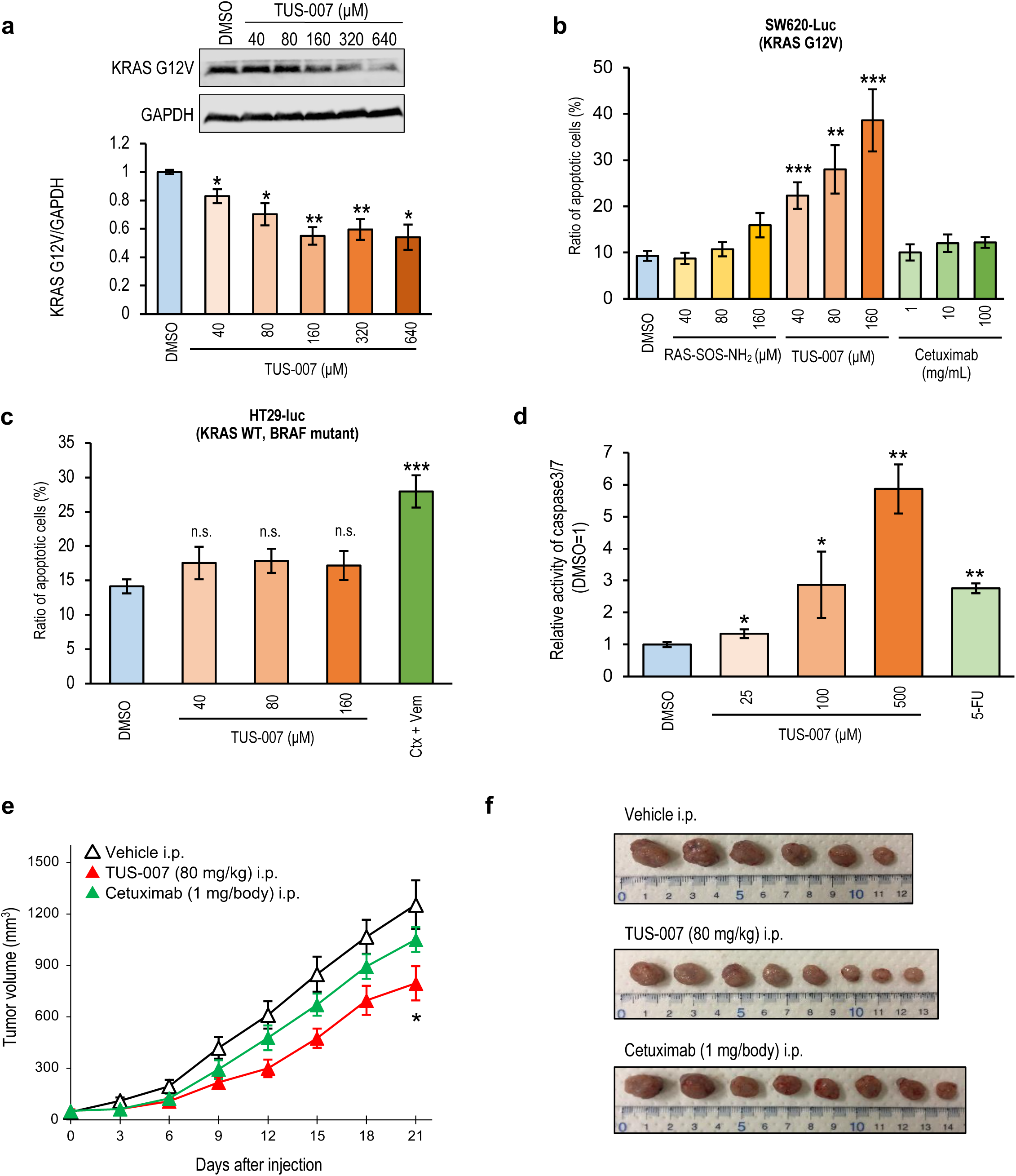
Chemical knockdown and tumor suppression *in vitro* and *in vivo* in KRAS G12V-driven human colon cancer cells by TUS-007 treatment. **a,** Immunoblotting analysis of KRAS G12V chemical knockdown in SW620-Luc cells treated with TUS-007 for 48 h at the indicated doses. The KRAS band intensity was normalized to that of GAPDH (mean ± SEM; n = 3). *P < 0.05 and **P < 0.01 vs. DMSO. **b & c,** The ratio of apoptotic cells were analyzed in colon cancer SW620-Luc cells (**b**) and HT29-Luc cells (**c**), a KRAS-independent, BRAF-dependent cell line (Ctx + Vem: 100 μg/ml cetuximab and 40 μM vemurafenib). Cells were treated with each compound for 48 h in terms of the number of annexin V-positive/PI-negative apoptotic cells detected by flow cytometry (mean ± SEM; n = 2–5). **P < 0.01, and ***P < 0.001 vs. DMSO. **d,** Relative activity of caspase 3/7 in SW620-Luc cells treated with TUS-007 or 5-fluorouracil (5-FU) for 48 h (mean ± SEM; n = 3). *P < 0.05 and **P < 0.01 vs. DMSO. **e,** Tumor volume in mice with SW620-Luc cells transplanted subcutaneously and treated with i.p. administration every 3 days (mean ± SEM; n = 6–8). *P < 0.05 vs. vehicle alone. **f,** Tumors of SW620-Luc cells at day 21 from the same mice shown in **e** (n = 6–8).

Next, to assess the effectiveness of TUS-007 *in vivo*, we transplanted SW620-Luc cells subcutaneously in immunodeficient mice. TUS-007 or cetuximab was administered to the xenograft mice by intraperitoneal (i.p.) injection. TUS-007 significantly attenuated tumor progression (Fig. 3e, f and Fig. S8a) and induced KRAS G12V chemical knockdown in tumors (Fig. S8b). The body weights of the mice were not affected by TUS-007 treatment (Fig. S8c). These results suggested that TUS-007 is effective *in vivo* against KRAS G12V-driven tumors.

### The effects of TUS-007 on KRAS G12D-driven pancreatic cancer

Approximately 95% of pancreatic cancers harbor KRAS mutations, and the G12D mutation is particularly prevalent and strongly correlated with poor prognosis (8). Therefore, we examined the effects of TUS-007 in human pancreatic cancer cell lines. As expected, TUS-007 degraded the KRAS protein (Fig. 4a) and induced apoptosis (Fig. 4b) in SW1990 KRAS G12D-driven pancreatic cancer cells (24). The apoptosis induction in SW1990 cells was approximately correlated with concentrations of chemical knockdown of KRAS (Fig. 4a and b). The correlation between the concentration of apoptosis induction and degradation induction was further confirmed (Fig. S9). In addition, apoptosis was induced with caspase 3/7 activation in SW1990 cells (Fig. 4c), which was counteracted by a proteasome inhibitor MLN but not by a ubiquitination inhibitor NAEi (Fig. 4d). Therefore, it suggested that TUS-007 induced apoptosis owing to target degradation independent of ubiquitination.

**Fig. 4.**
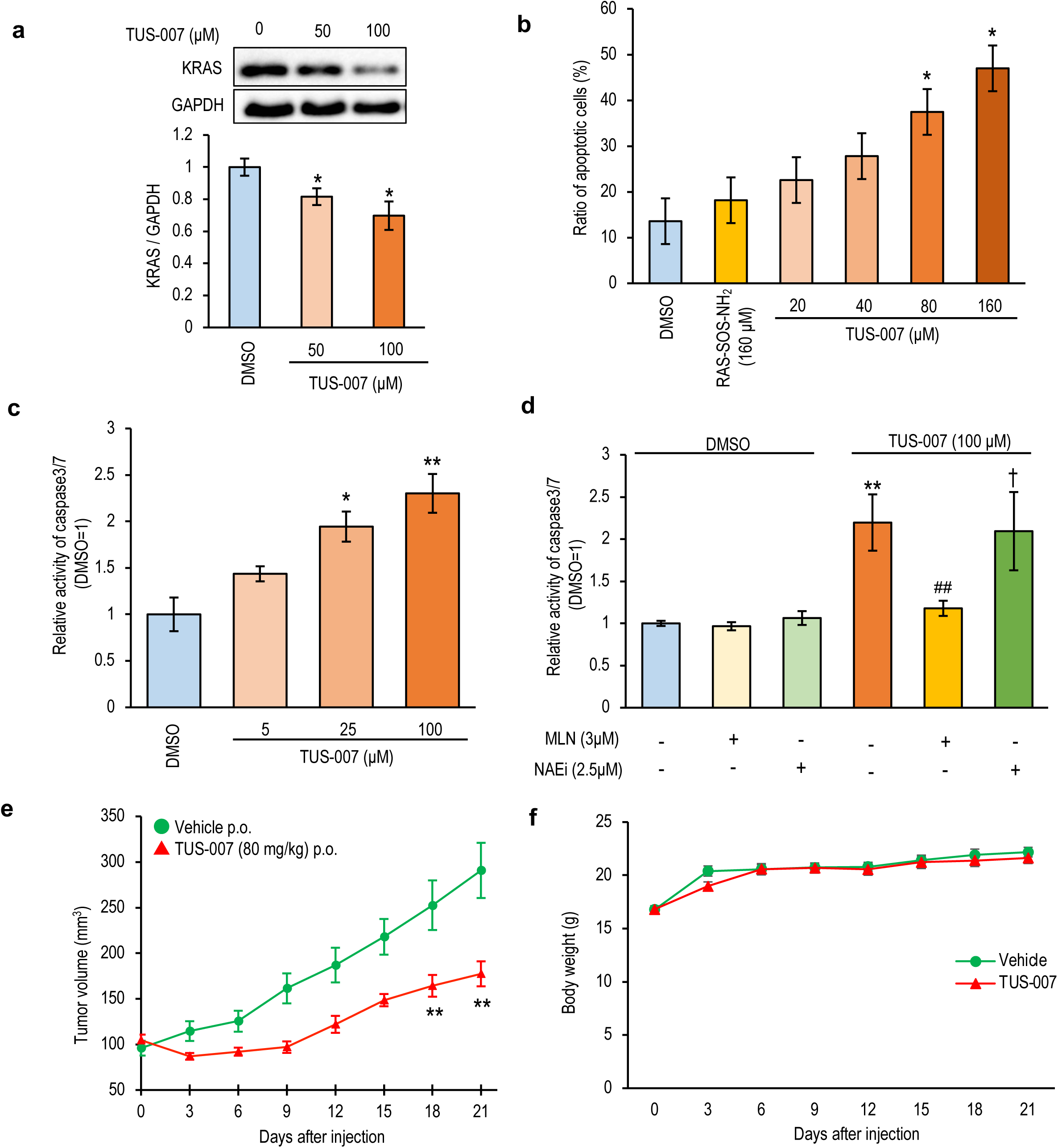
Chemical-knockdown and tumor suppression *in vitro* and *in vivo* in KRAS G12D-driven human pancreatic cancer cells by TUS-007 treatment. **a,** KRAS G12D chemical knockdown induced by TUS-007 in SW1990 cells. SW1990 cells were treated with TUS-007 for 72 h and analyzed by immunoblotting. The KRAS G12D band intensity was normalized to that of GAPDH (mean ± SEM; n = 4). *P < 0.05 vs. DMSO. **b,** The proportion of annexin V-positive apoptotic SW1990 cells treated with DMSO, RAS-SOS-NH_2_ or TUS-007 for 72 h (mean ± SEM; n = 3). *P < 0.05 vs. DMSO or RAS-SOS-NH_2_. **c,** TUS-007 increased the caspase 3/7 activity in SW1990 cells after treatment for 96 h (mean ± SEM; n = 3). *P < 0.05 and **P < 0.01 vs. DMSO. **d,** Caspase 3/7 activation by TUS-007 was proteasome dependent and ubiquitination independent. SW1990 cells were treated with the agents for 96 h (mean ± SEM; n = 3). **P < 0.01 vs. DMSO, ^##^P < 0.01 vs. TUS-007 (100 µM) only, and ^†^no statistically significant difference compared with TUS-007 (100 µM) only. **e,** Tumor volume in mice with SW1990 cells transplanted subcutaneously and treated with p.o. administered TUS-007 or vehicle (mean ± SEM; n = 6–9). The agents were administered every 3 days. **P < 0.01 vs. vehicle alone. **f,** Body weight changes in mice with SW1990 cells transplanted subcutaneously (mean ± SEM; n = 6–9).

In mice with SW1990 cells implanted subcutaneously, the oral (per os; p.o.) administration of TUS-007 significantly suppressed the tumor growth (Fig. 4e) and reduced tumor weight (Fig. S10a). The i.p. injection of TUS-007 also suppressed subcutaneously implanted tumor growth in mice (Fig. S10b). Both p.o. (Fig. 4f) and i.p. (Fig. S10c) administrations showed no change in body weight. Here, we examined the chemical knockdown of KRAS, HRAS and NRAS with TUS-007 in pancreas from the identical mice used in Fig. 4e and f. In accordant with the results in RAS-less MEFs, TUS-007 induced the degradation of KRAS and HRAS but not NRAS (Fig. S11). Thus, there was no sign of toxicity such as body weight losses in contrast to a pan-RAS inhibitor (21).

Moreover, extracellular signal-regulate kinase (ERK) and protein kinase B (AKT) phosphorylation were decreased in tumor by both p.o. and i.p. administrations (Fig. S12a and b), indicating that both RAS-mitogen-activation protein (RAS-MAP) and RAS-phosphoinositide 3-kinase (RAS-PI3K) signaling were inhibited. It is difficult for inhibitors to inhibit both RAS-MAP and RAS-PI3K signaling, even using a clinically effective KRAS G12C inhibitor (6). Thus, the chemical knockdown of KRAS by CANDDY might offer a clinical benefit. Taken together, these results indicated that TUS-007 exerted *in vivo* antitumor activity *via* the chemical knockdown of KRAS G12D/V and suppression of KRAS signaling.

### TUS-007 exerted anti-tumor activity even in orthotopic xenograft model mice

SW1990-Luc cells were transplanted directly into the pancreas of mice as an orthotopic xenograft model. The mice were subsequently treated p.o. with TUS-007. Remarkably, *in vivo* imaging revealed reductions in tumor growth (Fig. 5a, b and Fig. S13a) and tumor weight (Fig. S13b). Results from immunohistochemical analysis (Fig. 5c left panels and Fig. S13c left bar graph) and immunoblotting (Fig. 5d) confirmed the chemical knockdown of KRAS G12D in the tumor tissues of mice treated with TUS-007. Furthermore, higher concentrations of positive cells were observed in the terminal deoxynucleotidyl transferase dUTP nick-end labeling (TUNEL) assay in the tumor tissues of TUS-007-treated mice, implying the induction of apoptosis in the TUS-007-treated tumors (Fig. 5c right panels and Fig. S13c right bar graph). The body weights of the orthotopic xenograft model mice were not affected by TUS-007 (Fig. S13d). Overall, these results demonstrated that oral treatment with TUS-007 induces KRAS G12D chemical knockdown and suppresses pancreatic tumor growth without the sign of weight loss and significant toxicity in the orthotopic xenograft model.

**Fig. 5.**
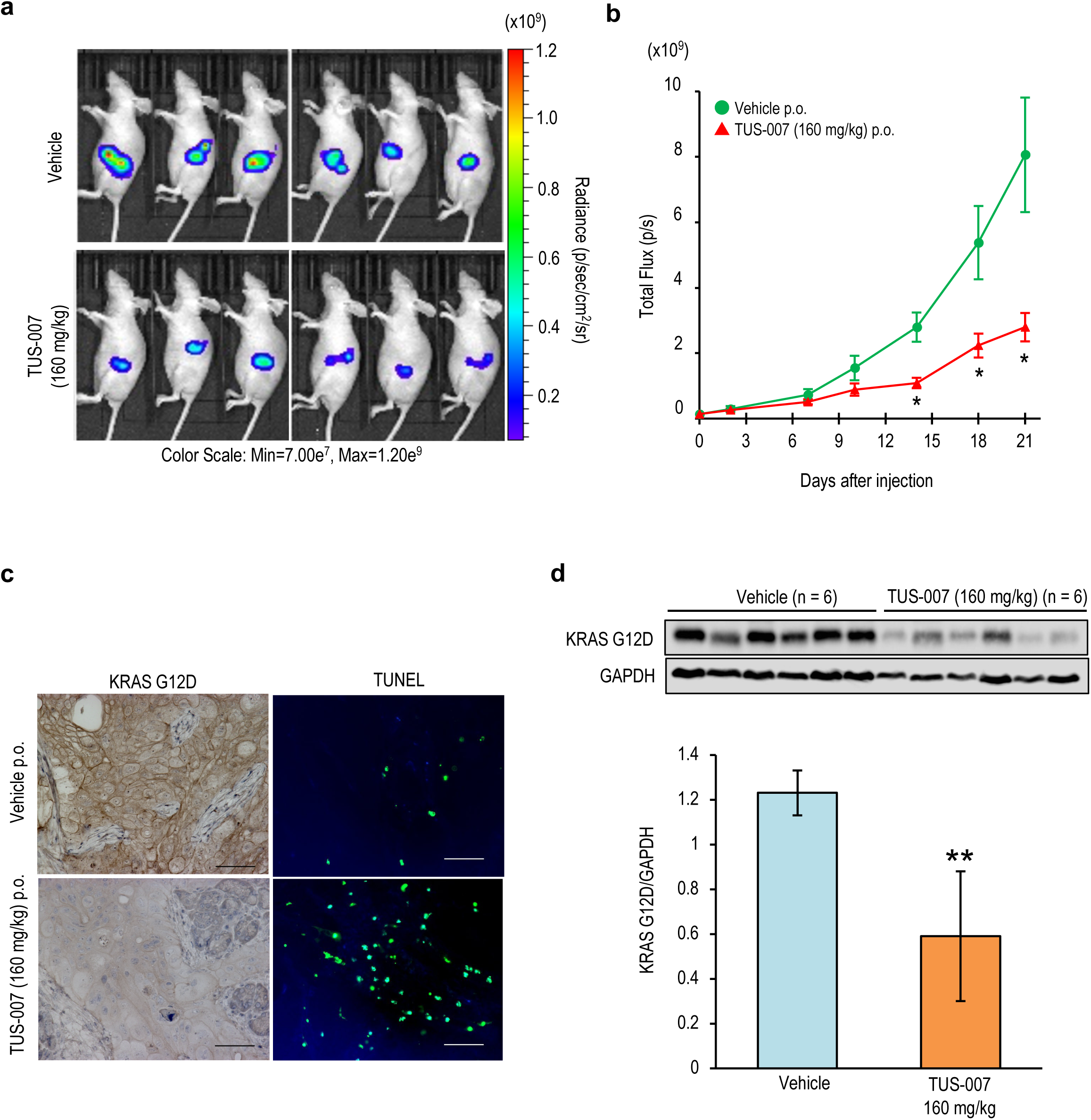
TUS-007 suppresses the growth of KRAS G12D-positive orthotopic pancreatic tumors. **a,** Images showing the luciferase activity and total flux in orthotopic tumors transplanted SW1990-Luc cells *in vivo*. **b,** Tumor growth (bioluminescence) in identical mice treated with vehicle or TUS-007 administered p.o. every 3 days. (mean ± SEM; n = 6). *P < 0.05 vs. vehicle alone. **c,** Representative histochemical staining of KRAS and TUNEL in orthotopic tumor sections from the same mice used in **a**. The scale bar is 60 µm. **d,** Immunoblotting of KRAS G12D in tumor lysates from the same mice used in **a**. The bar graph shows quantification of KRAS G12D normalized to GAPDH (mean ± SEM; n = 6). **P < 0.01 vs. vehicle alone.

We examined the *in vivo* concentrations of TUS-007. When an i.p dose of 80 mg/kg TUS-007 was administered to healthy mice, a concentration (45 ng/mg, equivalent to 53 µM) of TUS-007 was observed in the pancreas (Table S1.). This TUS-007 concentration *in vivo* agreed with the concentration at which chemical knockdown and apoptotic indicators were observed in SW1990 cells *in vitro* (Fig. 4a-c). Therefore, these results suggest that there is no contradiction between our *in vivo* and *in vitro* results.

Finally, to examine TUS-007 toxicity in detail, the general, histological, hematological and biochemical examinations using mice were performed. The p.o. treatment of TUS-007 at 80 mg/kg for continuous 5 days did not cause any toxic effect in body weight, organ weight, histology of organs, hematology and blood biochemistry at 2 weeks after the last treatment (Table S2-S6.). Thus, oral treatment with TUS-007 did not show sever toxicity.

### CANDDY applied to another undruggable target

To evaluate the versatility of CANDDY approach, we designed a CANDDY molecule targeting MDM2 (MDM2-CANDDY), a negative regulator of P53, using a P53-MDM2 PPI inhibitor with a weak potency (25) as an interactor module of a CANDDY molecule (Fig. S14a and Text S1.). MDM2-CANDDY successfully induced the chemical knockdown of MDM2 in HCT-116 human colon cancer cells (Fig. S14b). P53-MDM2 PPI is still well-recognized “undruggable” target (26). This result suggested the potential versatility of CANDDY to target other undruggable proteins.

## Discussion

We provided the proof of concept that CANDDY tag enabled to induce proteasomal degradation through ubiquitination-independent process (Fig. 1f and Fig. 2a) in this report. Considering the proteasome enable to degrade any targets, the CANDDY tag modified from a proteasome inhibitor (Fig. 1b) can eliminate the difficulties of matchmaking of targets and specific ubiquitination related proteins such as E3 ligases in current protein degradation (13, 14). Thus, CANDDY technology could provide a simple and rational approach to induce the chemical knockdown of currently undruggable targets because of the CANDDY tag independent of ubiquitination.

Expectedly, we confirmed the correlations between degradation of KRAS G12D and V and apoptosis in pancreatic and colon cancer cells, respectively (Fig. 3a, b, Fig. 4a-d, Fig. S7 and S9), and *in vivo* (Fig. 5c and Fig. S13c). Both *in vitro* and *in vivo*, TUS-007 achieved efficacy mediated by moderate degradations of KRAS G12D/V. A previous study showed that the moderate decrease of KRAS protein is enough to inhibit tumor growth *via* more strong inhibition of downstream signal (12) as well as we showed (Fig. S12). Thus, *in vitro* and *in vivo* results could be reasonable. Importantly, the concentration of TUS-007 in pancreas agreed with the effective concentration *in vitro* (Table S1.), demonstrating the consistency between our *in vivo* and *in vitro* results. On the other hand, the requirement of high concentrations generally raises concerns about toxicity. However, TUS-007 did not show non-specific cytotoxicity *in vitro* even at high concentration (Fig. 2c) and no sign of body weight losses was observed in TUS-007 treated mice (Fig. 4f, Fig. S8c, S10c and S13d) in contrast to a pan-RAS inhibitor (21). These results suggest that TUS-007 induced *in vivo* degradation of KRAS G12D/V resulting in suppression of colon and pancreatic tumors without no significant toxicity even at high concentrations.

No effective drug to inhibit or degrade KRAS G12D/V has been reported yet. Also, the proteolysis field does not still have enough evidence that can target and degrade tough and valuable undruggable proteins (14). Of note, in spite of a poor inhibitor was employed as interactor, TUS-007 induced chemical knockdown of KRAS G12D/V *in vivo*, resulting in suppression of a pancreatic tumor even in the orthotopic xenograft model with oral administrations (Fig. 5). In addition, another weak inhibitor of PPI, P53-MDM2 PPI inhibitor (25) (Fig. S14), was useful as the interactor of CANDDY molecules. Thus, CANDDY technology could be expectedly applied to tough and valuable undruggable targets still remaining with no effective inhibitors. In the future, CANDDY has the potential to overcome the current limitations of proteolysis and to expand the application scope of degraders for molecular targeting therapy of aggressive and deadly diseases caused by undruggable targets.

## Data availability

Source data are provided for all experiments. Other data that support the findings of this study are available from the corresponding author, upon reasonable request.

## Acknowledgments

We thank Masaaki Ozawa, Mai Hasegawa, Takuya Yogi, Mizuki Kanazawa, and Sachi Tsuji for help with the *in vitro* studies, Yuri Sato for help with the *in vivo* studies, Megumi Tsukamoto and Megumi Saito for help with both *in vitro* and *in vivo* studies. We thank the NCI RAS initiative for providing the RAS-less MEFs used in this work, the Uchiro Laboratory for performing the high-resolution mass spectrometry analysis at the Research Equipment Center (Tokyo University of Science (TUS), Japan), the Gunji Laboratory (TUS) for the use of their high-performance liquid chromatography instruments, the Iwakura and Abe Laboratories in the animal facility of the Research Institute for Biomedical Sciences (TUS) for maintaining the mice, Fumito Ishizuka and Naoki Takeda (NARD Institute, Ltd.) for contributions to the chemical synthesis, and Hiroyuki Ohashi for the helpful suggestions. This work was financially supported by the Program for Creating STart-ups from Advanced Research and Technology (START Program: 962238), and the joint research with FuturedMe Inc.

## Author contributions

S. Im., L. H., S. It., and E. M. S. designed the study. S. Im., L. H., S. It., M. I. and T. Y. performed the *in vitro* and *in vivo* experiments. Y. I. performed chemical analyses. E. M. S. supervised the study. All authors discussed the study and approved the submitted manuscript.

## Competing interests

The authors declare no competing interests.

**Supplementary information,** including Materials and methods, Text S1, Fig. S1-14 and Table S1-6, is available in the online version of the paper.

## Materials and methods

### Cell lines

SW1990 and HCT-116 cells were purchased from the American Type Culture Collection (Manassas, VA, USA). SW620-Luc and HT29-Luc cells were purchased from the National Institute of Biomedical Innovation, Health, and Nutrition (Osaka, Japan). RAS-less MEF cell lines (wild-type (wt) KRAS, KRAS G12D, KRAS G12V, KRAS G12C, NRAS and HRAS) were obtained from the National Institutes of Health (NCI RAS Initiative at the Frederick National Laboratory for Cancer Research, Frederick, MD, USA).

### Animal studies

BALB/cA-nu/nu and BALB/cA (female, 7-9 weeks old) were purchased from CLEA Japan (Tokyo, Japan). The animals were maintained in conventional housing conditions, with a daily 12-h light/dark cycle, and given food and water *ad libitum*. The animals were cared for in accordance with the NIH Guide for the Care and Use of Laboratory Animals. All animal experiments were approved by the Committee on Animal Experimentation of the Tokyo University of Science.

### Reagents

MLN2238 (A10600) was purchased from AdooQ BioScience (Irvine, CA, USA). NAEi was purchased from AdipoGen (San Diego, CA, USA). Human 26S Proteasome Protein (E-365) and Human 20S Proteasome Protein (E-360) were purchased from R&D Systems (Minneapolis, MN, USA). Human KRAS (G12D) and the corresponding His tag (12259-H07E1) were purchased from Sino Biological (Beijing, China). Human KRAS (G12V), 2-186, and the corresponding His tag (R06-32BH) were obtained from SignalChem Lifesciences (Richmond, British Columbia, Canada).

### Measurement of 20S proteasome activity

The chymotrypsin-like (β5), trypsin-like (β2), and caspase-like (β1) activities of the 20S proteasome were measured using a 20S Proteasome Activity Kit GOLD (StressMarq Bioscience Inc., Victoria, British Columbia, Canada). Purified 20S proteasome (0.1 µg) was incubated in the presence of CANDDY molecules (1, 4, 10, 20, 40, 80, and 160 µM) or MLN2238 at the indicated concentrations for 30 min at room temperature. After incubation, 100 mM of fluorogenic peptide substrates, Suc-LLVY-AMC (β5 substrate), Bz-VGR-AMC (β2 substrate), or Z-LLE-AMC (β1 substrate), was added, followed by incubation for 1 h at room temperature. The reaction mixture was then transferred to a 96-well plate, and the fluorecence from hydrolyzed AMC groups was measured using a Synergy HT multi-channel microplate reader (BioTek Instruments, Inc., Winooski, VT, USA) with a 360 nm excitation filter and 460 nm emission filter.

### Evaluation of affinity of TUS-007 for KRAS in a thermal shift assay

The KRAS G12D/V (100 nM) was preincubated with TUS-007, a RAS-SOS inhibitor, or 10% DMSO in 75 mM phosphate buffer (pH 7.5) for 30 min at 37 °C (G12D) or for 20 min at 25 °C (G12V). Aliquots of the reaction solution were sampled into separate microtubes and heated for 30 min at 70, 80, or 90 °C for G12D, or at 40, 50, or 60 °C for G12V. After centrifugation at 18,000 ×*g* for 20 min, the supernatants were analyzed by SDS-PAGE, followed by immunoblotting using an anti-KRAS antibody (1:1000, WH0003845MI, Sigma-Aldrich, St. Louis, MO, USA). Immunoreactive bands were detected using an iBright CL1000 chemiluminator (Thermo Fisher Scientific, Waltham, MA, USA) with an enhanced chemiluminescent substrate for detection of horseradish peroxidase (HRP; Merck, Darmstadt, Germany).

### Estimation of T_m_ values using fluorescence based thermal shift assay

To estimate the T_m_ values, we performed a fluorescence based thermal shift assay using ProteoStat Thermal Shift Stability assay (ENZO Life Sciences, Farmingdale, NY, USA). 1 µg of KRAS G12D was mixed with DMSO (2.5 %), RAS-SOS inhibitor (4 µM) or TUS-007 (4 µM) in 1x assay buffer containing a fluorescent indicator of protein destabilization, ProteoStat TS Detection Reagent, and incubated under heating from 25 °C to 99 °C in StepOne Plus Real-time PCR system (Thermo Fisher Scientific). The kinetics of fluorescent signals were analyzed to calculate T_m_ values in Protein Thermal Shift Software 1.3 (Thermo Fisher Scientific).

### Evaluation of KRAS degradation induced by TUS-007 in a cell-free system

KRAS G12D (final concentration: 5 ng/µL) and 26S proteasome (final concentration: 8 nM) were incubated with TUS-007 (2, 3, 5, 10, 20, 40 µM), RAS-SOS inhibitor (40 µM), RAS-SOS-NH_2_ (40 μM) or DMSO in 20 mM Tris-HCl buffer (pH 7.5) containing 20% glycerol for 3 h at 37 °C. To assess the effect of MLN on chemical knockdown, KRAS G12D and 26S proteasome were incubated with DMSO or TUS-007 40 µM in the presence or absence of MLN (1 µM) in the same condition. KRAS G12V (final concentration: 5 ng/µL) and 26S proteasome (final concentration: 8 nM) were incubated with TUS-007 at the indicated concentration or DMSO in 75 mM phosphate buffer, containing 1% LABRASOL (Gattefossé, Saint-Priest, France), for 1 h at 37 °C. After centrifugation at 14,000 ×*g* for 10 min, the supernatants were analyzed by SDS-PAGE, followed by immunoblotting using an anti-KRAS antibody (1:1000, WH0003845MI for G12D, Sigma-Aldrich; and 1:2000, F234 for G12V, Santa Cruz Biotechnology, Inc., Dallas, TX, USA). The immunoreactive bands were detected, as described above. The DC_50_ values were calculated by a Rodbard approximation using the band intensity obtained from four and three independent experiments for G12D. Immunoblotting was performed as mentioned above.

### Evaluation of target degradation in HeLa cells by flow cytometry

The expression plasmid pMIR-DsRed-IRES-ecDHFR-HA-GFP was a kind gift from Dr. L. Hedstrom (Brandeis University). Plasmid expressing human KRAS G12D was constructed by RF cloning with some minor modifications. A full-length cDNA clone of human KRAS (AK292510) was obtained from the National Institute of Technology and Evaluation (Tokyo, Japan). Hybrid RF cloning primers were designed using RF-Cloning.org. All PCR amplifications were performed with KOD-Plus-Neo (Toyobo Co. Ltd., Japan). The plasmid DNA sequences encoding target proteins were determined using an ABI 3500 (Applied Biosystems, MA, USA). HA-GFP-fusion target protein- and DsRed-expressing HeLa cells were treated with 1% DMSO or CANDDY molecules in serum-free medium for 24 h. The cells were harvested using trypsin and resuspended in fluorescence activated cell sorter (FACS) buffer. The cells were then analysed on a BD FACSCanto II. Data were analysed using FlowJo software (FlowJo LLC).

### Cell proliferation assay

To examine the effect of TUS-007 on cell proliferation, the WST-8 assay (Cell Count Reagent SF; Nacalai Tesque, Kyoto, Japan) was performed. RAS-less MEF cells, expressing human RAS, were plated in 96-well plates at a density of 3 × 10^3^ cells per well, cultured overnight, then treated with the indicated concentrations of TUS-007 in 4% FBS containing medium (KRAS G12D), 10% FBS containing medium (KRAS G12V) or serum-free medium (KRAS G12C, wt KRAS, HRAS, NRAS) for 72 h. After incubation, cell viability was measured spectrophotometrically using the WST-8 reagent. The culture medium was removed and WST-8 reagent was mixed with the growth medium at a ratio of 1:10 (with a final volume of 110 µL) and added to each well. The cells were incubated in an atmosphere of 5% CO_2_ at 37 °C for 1 to 2 h, and the absorbance at 450 nm was measured using the aforementioned Synergy HT multi-channel microplate reader (BioTek Instruments, Inc.).

### Evaluation of TUS-007 specificity in cells

RAS-less MEFs were used to evaluate the selective degradation by TUS-007. RAS-less MEFs expressing one of the human WT RAS members (KRAS WT, HRAS WT, or NRAS WT) or KRAS G12C were plated into 6-well plates at a density of 3 × 10^4^ cells per well, cultured overnight, and treated with the indicated concentrations of TUS-007 in serum-free medium for 72 h. RAS-less MEFs expressing KRAS G12D or G12V were plated into 96-well plates at a density of 3 × 10^3^ cells per well, cultured overnight, and treated with the indicated concentrations of TUS-007 in 4% FBS or 10% FBS containing medium, respectively, for 72 h. The cells were lysed using RIPA buffer (Nacalai Tesque., Kyoto, Japan). The lysates were centrifuged at 18,000 ×*g* for 10 min. The supernatants were analyzed by western blotting using mouse monoclonal antibodies specific to KRAS (1:1000, WH0003845MI; Sigma-Aldrich) and glyceraldehyde 3-phosphate dehydrogenase (GAPDH, 1:10000, sc-32233; Santa Cruz Biotechnology, Inc.) and rabbit polyclonal antibodies specific for HRAS (1:1000, 18295-1-AP; Proteintech, Rosemont, IL, USA) and NRAS (1:1000, 10724-1-AP, Proteintech). Immunoreactive bands were detected using iBright CL1000 chemiluminator or LAS 3000 (Fujifilm, Tokyo, Japan) with an enhanced chemiluminescent substrate for the detection of horseradish peroxidase (HRP; Merck, Darmstadt, Germany). The intensity of the RAS band was quantified using ImageJ software and normalized to the intensity of the GAPDH band.

### Measurement of KRAS protein levels in cells by western blotting

SW620-Luc and SW1990 cells were treated with 1% DMSO or TUS-007 in serum-free medium for 12 or 48 h (SW620-Luc cells) and 12 or 72 h (SW1990 cells). The cells were lysed using a RIPA buffer (Nacalai Tesque., Kyoto, Japan). The precipitates were separated from the soluble fraction by centrifugation at 18,000 ×*g* for 20 min. The supernatants were analyzed by western blotting with mouse monoclonal antibodies specific for KRAS (1:1000, WH0003845MI; Sigma-Aldrich) and GAPDH (1:20000, sc-32233; Santa Cruz Biotechnology, Inc.). The immunoreactive bands were detected by LAS-3000, as described above. The intensity of the KRAS band was quantified using ImageJ software and normalized to the intensity of the GAPDH band.

### Evaluation of caspase activity

To measure caspase activity, SW1990 cells were plated onto 96-well white plates and 96-well clear plates, at a concentration of 8 × 10^3^ cells/well, in a medium containing 10% FBS. The plates were incubated overnight without CO_2_ equilibration at 37 °C. The medium was replaced with medium containing 2% FBS, DMSO, 100 µM TUS-007, 3 µM MLN2238, or 2.5 µM NAEi and incubated for 96 h. SW620-Luc cells were plated onto 96-well white plates and 96-well clear plates at 2 × 10^3^ cells/well in medium containing 10% FBS. After overnight incubation, the medium was replaced with FBS-free medium containing 1% L-glutamine, DMSO, and 25, 100, or 500 µM TUS-007 for 48 h. The caspase 3/7 Glo assay (Promega, Madison, WI, USA) was performed on the white plates according to the manufacturer’s protocol. The chemiluminescence intensity was measured with a Synergy HT plate reader (BioTek). The cell viability was measured in the clear plates using the WST-8 assay, as described above. Caspase 3/7 activity was obtained as the chemical luminescence intensity of the caspase 3/7 Glo assay normalized to the cell viability.

### Analysis of apoptosis by flow cytometry

SW1990, SW620-Luc, and HT-29 cells were seeded in 24-well plates, followed by the addition of compounds at the indicated concentrations. After incubation for 12 or 48 h (SW620-Luc, HT-29) and 12 or 72 h (SW1990), the cells were harvested using trypsin, washed with phosphate-buffered saline (PBS), and pelleted by centrifugation at 110 ×*g* for 5 min. The cells were resuspended in 85 µL of binding buffer (Annexin V-FITC Kit, Medical & Biological Laboratories Co., Ltd., Nagoya, Japan). Subsequently, 10 µL of annexin V-FITC and 5 μL of propidium iodide (PI) were added to each sample, followed by incubation in the dark at room temperature for 15 min. After incubation, 400 µL of binding buffer was added to each sample on ice. The cells were then analyzed using a FACSCanto II fluorescence-activated cell sorter (BD Biosciences, Santa Clara, CA, USA) to detect the fluorescein isothiocyanate (FITC) signal with excitation and emission filter wavelengths of 488 nm and 580 nm, respectively, to detect the PI signal. Data were analyzed using FlowJo software (FlowJo, Eugene, OR, USA).

### Subcutaneous xenograft model

SW620-Luc (3 × 10^7^ cells/mL) and SW1990 (5 × 10^7^ cells/mL) cells were suspended in PBS, and 100 µL of the single-cell suspension was transplanted subcutaneously into the right flanks of BALB/cA-nu/nu mice using a 26 G syringe. Tumor volumes were calculated as the tumor length × width^2^ × 0.5. When the tumor volumes reached approximately 100 mm^3^, the mice were randomized into treatment and control groups of 6 to 8 animals per group. For i.p. treatment, TUS-007 was dissolved in DMSO and then diluted to a final concentration of 10% DMSO in 20% polyethylene glycol (PEG) 400/Tween 80 (1:1 ratio). The TUS-007 solution was administered intraperitoneally into the tumor-bearing mice at a dose of 80 mg/kg every three days. The control mice were treated with the PEG/Tween vehicle alone. For p.o. treatment, TUS-007 was suspended in 0.5% (w/v) carboxymethyl cellulose (CMC) and administered p.o. at a dose of 80 mg/kg every three days. CMC (0.5% w/v) was administered to control mice. Cetuximab was administered to an additional group of SW620-Luc-implanted mice at a dose of 1 mg/mouse every three days.

### Orthotopic xenograft model

The pGL4.51 (Luc2/CMV/Neo) vector (Promega) was transfected into SW1990 cells. Successfully transfected cells were selected using G418. SW1990-Luc cells were mixed with growth factor-reduced Matrigel (BD Biosciences) on ice, at a ratio of 1:1 (v/v), to obtain a final cell density of 1.5 × 10^4^ cells/µL. Ten microliters of the cell suspension was injected directly into the pancreas of BALB/cA-nu/nu mice using a 27 G syringe. Three days after cell transplantation, TUS-007 treatment was initiated in six mice. TUS-007 was suspended in 0.5% (w/v) CMC and administered orally at a dose of 160 mg/kg every 3 days. CMC (0.5% w/v) was administered to control mice (n = 6). Tumor luciferase activity was monitored based on the bioluminescence intensity. Fifty microliters of D-luciferin (30 mg/mL in saline) was injected subcutaneously in mice (Promega). Twenty minutes after the injection, bioluminescence images were obtained using an IVIS Lumina LT (Perkin Elmer, Waltham, MA, USA) with an exposure time of 15 s while the mice were under isoflurane anesthesia (Wako, Tokyo, Japan).

### Measurement of RAS levels and downstream signaling molecule activation in xenograft tumors

KRAS expression in xenograft tumors and wt RAS protein expression in pancreas were evaluated by immunohistochemistry and western blot analysis. Tumor tissue and pancreas were harvested from the mice harboring subcutaneous xenografts on day 21 of the TUS-007 administration. Tumor tissues were harvested from the orthotopic xenograft mice on day 24 after the TUS-007 administration for consecutive 3 days starting on day 21. A portion of each sample was fixed in 4% paraformaldehyde (Nacalai Tesque., Kyoto, Japan), embedded in paraffin, then cut into 5 µm-thick sections. The sections were deparaffinized and incubated in 0.3% hydrogen peroxide in methanol for 30 min, and antigen retrieval was performed using a 10 mM citrate buffer (pH 6.0) at 121 °C for 20 min. Tumor tissues were subsequently blocked with 1% bovine serum albumin (BSA) in PBS for 30 min and stained with primary antibodies specific for human KRAS (1:200, 12063-1-AP; Proteintech), diluted in 1% BSA, followed by HRP-labeled secondary antibodies (1:600, ab6721; Abcam, Cambridge, UK) in 1% BSA for 30 min at room temperature. The sections were counterstained with hematoxylin and visualized with diaminobenzidine (1.02924.0001; Merck). Digital images were obtained at 40× magnification using a model BZ-9000 microscope (Keyence, Osaka, Japan). The area of KRAS staining was quantified using ImageJ software, and the perimeter-to-area ratio (KRAS-positive area/section) was calculated for a total of 30 images of 6 tumor sections from each group. The remaining part of each tumor sample was homogenized in a cell-lysis buffer (50 mM triethanolamine, 50 mM KCl, 5 mM MgCl_2_, 0.25 M sucrose, 1 mM phenylmethylsulfonyl fluoride, proteinase inhibitors (Nacalai Tesque., Kyoto, Japan), 1 mM dithiothreitol, and ribonuclease inhibitor (0.2 unit/µl, TaKaRa Bio., Shiga, Japan) for western blot analysis, as described above. Mouse monoclonal antibodies specific for KRAS (1:1000, WH0003845MI; Sigma-Aldrich), HRAS (1:1000, 18295-1-AP; GAPDH (1:20000, sc-32233; Santa Cruz Biotechnology, Inc.) and rabbit antibodies specific for p-Erk1/2 (phospho-p44/42 MAPK) (1:1000, #4370; Cell Signaling Technology, Beverly, MA, USA), Erk1/2 (p44/42 MAPK) (1:1000, #4695; Cell Signaling Technology), p-Akt (1:2000, #4060; Cell Signaling Technology), and Akt (1:2000; #4691; Cell Signaling Technology) were used for the immunohistochemical staining. The immunoreactive bands were detected by LAS-3000 as described above. Quantitative analysis of the western blots was performed using ImageJ software, and the results were normalized to the GAPDH band intensity.

### Measurement of apoptosis in xenograft tumors

DNA fragmentation in the tumor tissues was evaluated by a TdT-mediated dUTP nick-end labeling (TUNEL) assay. Paraffin-embedded tissue sections (5 µm thick) were deparaffinized. The TUNEL reaction was performed using the DeadEnd Fluorometric TUNEL System (Promega), according to the manufacturer’s instructions. Fluorescence images were acquired using a BZ-9000 fluorescence microscope (Keyence). TUNEL-positive cells were identified and counted using ImageJ software. In total, 30 images of 6 tumor sections were analyzed from each group.

### Determination of TUS-007 in pancreas

BALB/cA mice were treated i.p. with TUS-007, and pancreas tissue was collected at suitable time points. Approximately 30-60 mg of tissue was homogenized in 0.6 mL of tissue-lysis buffer (50 mM Tris-HCl pH 8.0, 20 mM EDTA, 10 mM NaCl, 1% SDS) by Micro Smash MS-100 using stainless (5.5 φ) and zirconia (1.0 φ) beads (TOMY SEIKO CO., LTD., Tokyo, Japan), and 1 μL propyl p-hydroxybenzoate (4 mg/mL in DMSO) was added to each sample as an internal standard. Lysates were centrifuged at 18,000 ×*g* for 10 min and 2.25 mL of methanol/chloroform (ratio of 2:1) was added to the supernatants and the mixture was allowed to stand for 30 min at room temperature. Chloroform and distilled water (0.75 mL each) were added and samples were centrifuged at 1,500 ×*g* for 15 min. The layer of chloroform was collected. The supernatants were dried under reduced pressure and redissolved in the mobile phase. TUS-007 was determined by HPLC (Alliance e2695, Waters Corporation., MA, USA) using COSMOSIL^®^ C18-AR-II column (150×4.6 mm, Nacalai Tesque., Kyoto, Japan) in the isocratic elution mode with 0.1% trifluoroacetic acid (TFA) in 50:50 (v/v) acetonitrile/water mobile phase at a flow rate of 1.0 mL/min. All reagents for the assay were purchased from Nacalai Tesque. (Kyoto, Japan).

### Toxicity tests of TUS-007 in mice

The male ICR mice (6 weeks old) were administrated vehicle solution (0.5% CMC containing 2.5% DMSO) or TUS-007 dissolved vehicle at 80 mg/kg for continuous 5 days. The observation of general conditions was performed every day up to Day 17. The body weight was measured at Day 0, 1, 2, 3, 4, 6, 8, 10, 12, 15 and 17. At Day 17, the organs and blood were collected under isoflurane anesthesia. The organs were immediately fixed in 10% phosphate buffered formalin and embedded in paraffin for histological examination after HE staining. The blood cells were analyzed by FACS (ADVIA2120i, Siemens Healthineers AG, Germany.) and the blood biochemical examinations were performed using automatic analyzer TBA-120FR (CANON MEDICAL SYSTEMS Co., Ltd., Tochigi, Japan). Statistical analyses were performed with SAS Release 9.1.3 (SAS Institute Inc., NC, USA).

### Degradation of MDM2 in HCT-116 cells

HCT-116 human colon-cancer cells were treated with 1% DMSO or TUS-007 in serum-free medium for 24 h. The cells were lysed using a RIPA buffer (Nacalai Tesque). The precipitates were separated from the soluble fraction by centrifugation at 18,000 ×*g* for 20 min. The supernatants were analyzed by western blotting using a rabbit polyclonal antibody specific for MDM2 (1:1000, sc-965, Santa Cruz Biotechnology, Inc.) and GAPDH (1:20000, sc-32233; Santa Cruz Biotechnology, Inc.). The immunoreactive bands were detected by LAS-3000, as described above. The intensity of the MDM2 band was quantified using ImageJ software and normalized to the intensity of the GAPDH band.

### Statistical analyses

Statistical significance was determined with JMP^®^ Pro 14 (SAS Institute Inc., Cary, NC, USA). An unpaired *t-*test was used to compare pairs of groups under the assumption of normality. A one-way ANOVA with Dunnett’s post hoc analysis was used to compare sets of three or more groups. P-values < 0.05 were considered statistically significant.

## Text S1. Synthesis of CANDDY molecules

### General

Nuclear magnetic resonance (NMR) spectra were recorded on a Bruker BioSpin AVANCE600 spectrometer operating at 600 MHz for ^1^H-NMR. Chemical shifts were reported in the scale relative to CD_3_OD as an internal reference. Fast-atom bombardment mass spectrometry was performed on a JEOL JMS-700. Column chromatography was performed with silica gel 60 (75 mm) purchased from Nacalai Tesque, Inc. PLC silica gel 60 was purchased from Merck KGaA. Reversed phase liquid chromatography was performed with high performance liquid chromatography (HPLC) (Shimadzu Co., Ltd), with a YMC-Pack ODS-A or Inertsil ODS-4 (GL Science) (250 × 4.6 mm), an LC-6AD pump, and a RID-10A detector (Shimadzu Co., Ltd). HPLC experiment was performed with isocratic condition A (MeCN/H_2_O/TFA, 70:30:0.1; flow rate, 1 mL/min) for compounds **1**–**3**, and condition B (MeCN/H_2_O/TFA, 55:45:0.1; flow rate, 1 mL/min) for TUS-007.

H-Gly-OtBu · HCl and 2,5-dichlorobenzoic acid were purchased from Ark Pharm. H-Leu-OtBu · HCl and H-Phe-OtBu · HCl were purchased from Watanabe Chemical Industry.

Synthesis of TMP-NH_2_ was based on a previous report^26^. The synthesis of RAS-SOS inhibitor (**4**) and RAS-SOS-NH_2_ (**5**) was based on a previous report^15^. A part of the compound was synthesized by NARD Institute. Ltd. (Kobe, Japan).

### The synthesis of TUS-007

TUS-007 was synthesized as shown below.

Some of the compounds were synthesized by the NARD Institute. Ltd. (Kobe, Japan).

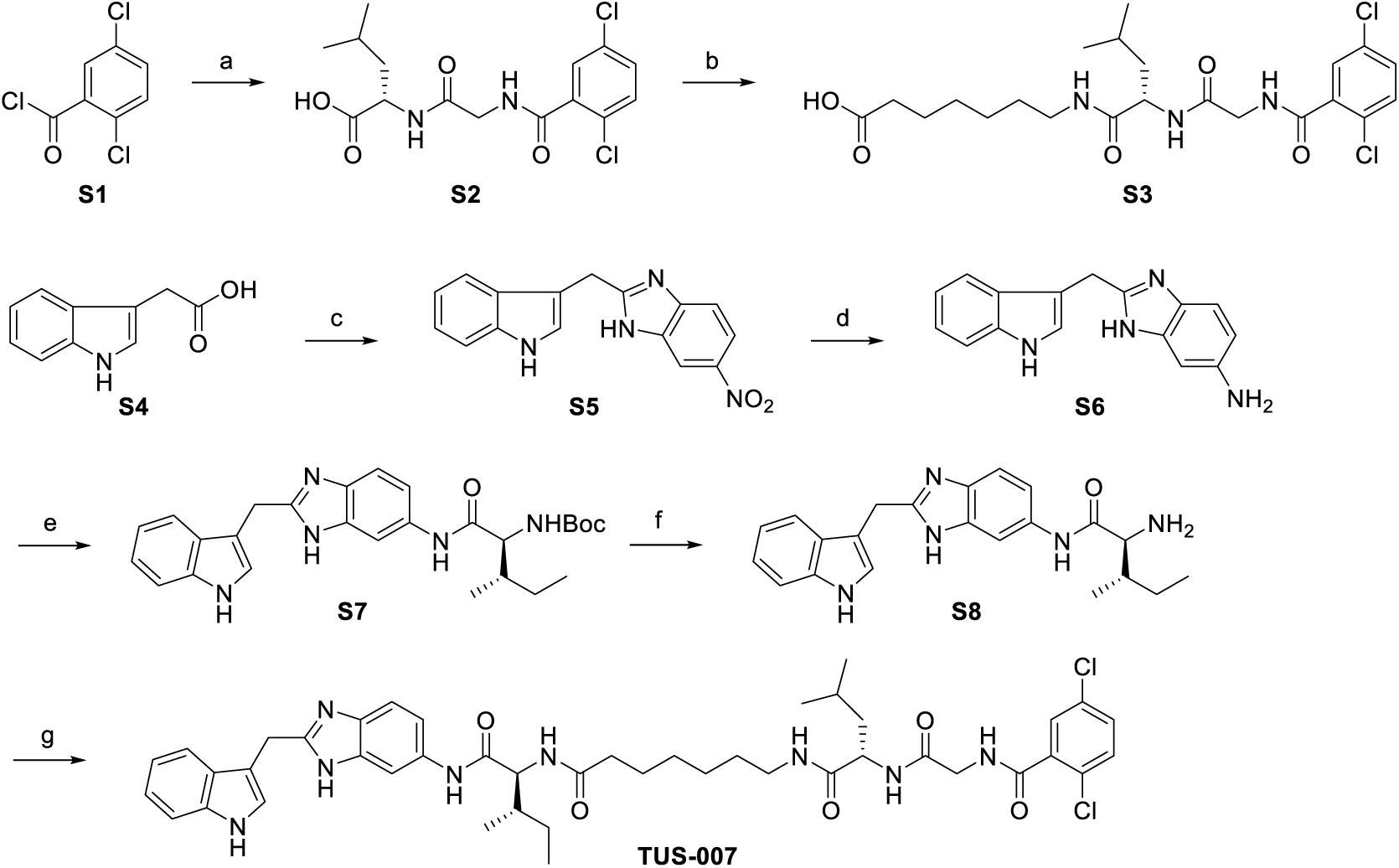
Reagent and conditions: (a) Glycyl-L-luecine, 1N NaOHaq.,THF 0°C, 88%; (b) (i) HOSu, EDC-HCI, CH_2_CI_2_, 0°C to rt, (ii) 7- Amino-heptanoic acid, Triethylamine, DMF, 0°C to rt, 72%; (c) (i) CDI, THF, 0°C to rt; 4-Nltro-1,2-phenylenedlamine, 47°C, (ii) AcOH, reflux, 88%; (d) H_2_, Pd/C, MeOH, rt, 62%; (e) *N*-Boc-L-lsoleucine, COMU, DIPEA, MeCN, 0°C to rt, *quant*.; (f) TFA, CH_2_CI_2_, rt, 76%; (g) **S3**, HATU, DIPEA, DMF, 0°C to rt, 53%.

To a solution of glycyl-L-leucine (125 g, 664 mmol) in 1N NaOH*aq*. (1.3 L) with tetrahydrofuran (THF: 550 mL), 2,5-dichlorobenzoyl chloride (127 g, 604 mmol) in THF (250 mL) was added dropwise at 0°C. After stirring for 2 h at the same temperature, the resulting solution was acidified (ca. pH2) with 2N HCl*aq*. and extracted with CHCl_3_ (total 3 L). The combined organic layer was washed with brine, dried over MgSO_4_, filtered, and concentrated *in vacuo*. The residue was suspended in diisopropyl ether (IPE: 1.2 L) and stirred for 30 min at 50°C. The resulting suspension was cooled to room temperature, and a precipitate was filtered, washed with IPE (500 mL), and thus **S2** (194 g, 88 % yield, white solid) was obtained.

i. To a solution of **S2** (194g, 536 mmol) in CH_2_Cl_2_ (1.6 L), *N*-hydroxysuccinimide (HOSu: 2.5 g, 804 mmol) and 1-ethyl-3-(3-dimethylaminopropyl) carbodiimide hydrochloride (EDC-HCl: 54 g, 804 mmol) were added at 0°C under Ar atmosphere. After stirring for 3 h at room temperature, the resulting solution was washed with water (1 L) three times, dried over MgSO_4_, filtered, and concentrated *in vacuo*. (ii) The residue (309 g) was dissolved in DMF (1.5 L) at Ar atmosphere, and 7-aminoheptanoic acid (77.8 g, 536 mmol) and triethylamine (178 mL) were added to the solution at 0°C. After stirring overnight at room temperature, 1N HCl*aq*. (1.5 L), water (1 L) and AcOEt (3 L) were added and extraction was performed using AcOEt (1.5 L). The combined organic layer was dried over Na_2_SO_4_, filtered, concentrated *in vacuo* and azeotroped with toluene. The residue (394 g) was purified by silica gel column chromatography (4.5 kg, elution with 0-5 % MeOH/CHCl_3_) and thus **S3** (190 g, 72 % yield, white solid) was obtained. **S3**: ^1^H-NMR (400 MHz, DMSO-d_6_) δ 0.84 (3H, d, *J* = 6.4 Hz), 0.87 (3H, d, *J* = 6.4 Hz), 1.18-1.27 (4H, m), 1.35-1.60 (7H, m), 2.16 (2H, t, *J* = 7.3 Hz), 2.94-3.09 (2H, m), 3.82-3.92 (2H, m), 4.26-4.31 (1H, m), 7.53 (3H, s), 7.92 (1H, t, *J* = 5.5 Hz), 7.96 (1H, d, *J* = 8.2 Hz), 8.78 (1H, t, *J* = 6.0 Hz), 11.96 (1H, s).
ii. To a solution of indol-3-ylacetic acid (100 g, 571 mmol) in THF (200 mL), 1,1’-Carbonyldiimidazole (CDI: 109 g, 674 mmol) was added in portions over 15 min at 0°C under Ar atmosphere. After stirring for 30 min at room temperature, 4-nitro-1,2-phenylenediamine (96.2 g, 628 mmol) was added, and stirred for 30 min at same temperature. The resulting solution was warmed to 47°C and concentrated *in vacuo* after stirring overnight. (ii) The residue was dissolved in AcOH (2 L) with ice-cooling and heated to reflux for 2 h. The resulting solution was concentrated *in vacuo* and azeotroped with toluene. The residue was dissolved in AcOEt (1 L), and insoluble portion was filtered on a Celite-pad, then washed with AcOEt (0.8 L). The filtrate was washed with water (1.5 L) and brine (1 L), dried over Na_2_SO_4_, and concentrated *in vacuo*. The residue (298 g) was purified by silica gel column chromatography (2 kg, elution with 0-30 % AcOEt/CHCl_3_) and thus **S5** (146 g, 88 % yield, brown amorphous) was obtained.

To a solution of **S5** (146 g, 500 mmol) in MeOH (1 L), Pd/C (20.0 g) was added and stirred overnight at room temperature under H_2_ atmosphere. The resulting solution was filtered on a Celite-pad, washed with MeOH (2 L), and concentrated *in vacuo*. The residue was purified by silica gel column chromatography (3.0 kg, elution with 5-20 % MeOH/CHCl_3_) and thus **S6** (85.4 g, 62 % yield, black solid) was obtained.

To a solution of **S6** (81.9 g, 351 mmol) in MeCN (1.2 L), *N*-ethyldiisopropylamine (DIPEA: 119 mL) and (1-Cyano-2-ethoxy-2-oxoethylidenaminooxy)dimethylamino-morpholinocarbenium Hexafluorophosphate (COMU: 58 g, 368 mmol) were added at 0°C under Ar atmosphere, and stirred for 1 h at room temperature. After cooling to 0°C, *N*-Boc-isoleucine (89.4 g, 341 mmol) was added and stirred overnight at room temperature, and then concentrated *in vacuo*. The residue was diluted with CHCl_3_ (2 L) and washed with water (1.5 L) twice, concentrated *in vacuo*, and thus **S7** (330 g, *quant.*, black oil) was obtained. This compound was used in the next step without further purification.

To a solution of **S7** (330 g) in CH_2_Cl_2_ (1.6 L), trifluoroacetic acid (TFA: 800 mL) was added at room temperature under Ar atmosphere and stirred for 2 h at the same temperature. The resulting solution was concentrated *in vacuo*, and then dissolved in AcOEt (1 L), washed with *sat*. NaHCO_3_*aq*. (600 mL) twice, dried over Na_2_SO_4_, filtered, and concentrated *in vacuo*. The residue (315 g) was purified by NH-silica gel column chromatography (2.3 kg, elution with 0-5 % MeOH/CHCl_3_), and thus **S8** (97.5 g, 76 % yield, yellow solid) was obtained.

To a solution of **S3** (139 g, 285 mmol) in *N*,*N*-dimethylformamide (DMF: 1250 mL), DIPEA (77.1 mL) and 1-[bis(dimethylamino)methylene]-1*H*-1,2,3-triazolo[4,5-*b*]pyridinium 3-oxide hexafluorophosphate (HATU: 109 g, 259 mmol) were added at 0°C under Ar atmosphere. After stirring for 30 min at the same temperature, **S8** (97.4 g, 259 mmol) was added at 0°C and stirred overnight at room temperature. The resulting solution was diluted with AcOEt (2.5 L) and water (1.5 L) and extracted with AcOEt (1.5 L). The combined organic layer was concentrated *in vacuo* and azeotroped with toluene. The residue (448 g) was purified by NH-silica gel (6.0 kg, elution with 0-4 % MeOH/CHCl_3_) and concentrated *in vacuo*. After dissolving in MeOH (500 mL), insoluble was filtered off and the filtrate was concentrated *in vacuo*. The residue was suspended in Me_2_CO (1.5 L), stirred, and cooled to room temperature. Subsequently, a precipitate was collected, and thus **TUS-007** (117 g, 53 % yield, yellow solid) was obtained.**TUS-007**: ^1^H-NMR (400 MHz, DMSO-d_6_) δ 0.83 (3H, d, *J* = 6.4 Hz), 0.74-0.88 (6H, m), 0.87 (3H, d, *J* = 6.4 Hz), 1.08-1.28 (5H, m), 1.36-1.59 (8H, m), 1.75-1.82 (1H, m), 2.07-2.21 (2H, m), 2.94-3.08 (2H, m), 3.81-3.92 (2H, m), 4.22-4.38 (4H, m), 6.89-6.93 (1H, m), 7.02-7.06 (1H, m), 7.14-7.44 (4H, m), 7.34 (1H, d, *J* = 8.2 Hz), 7.52 (3H, s), 7.88-7.97 (4H, m), 8.78 (1H, t, *J* = 5.7 Hz), 9.96 (1H, d, *J* = 22.4 Hz), 10.92 (1H, s), 11.93-11.97 (1H, m). HR-MS (FAB) *m*/*z*: 845.3674 (Calcd for C_44_H_55_Cl_2_N_8_O_5_ [M+H]^+^ 845.3672).

### The synthesis of MDM2-CANDDY

MDM2-CANDDY was synthesized as shown below. Some of the compounds were synthesized by the NARD Institute. Ltd. (Kobe, Japan).

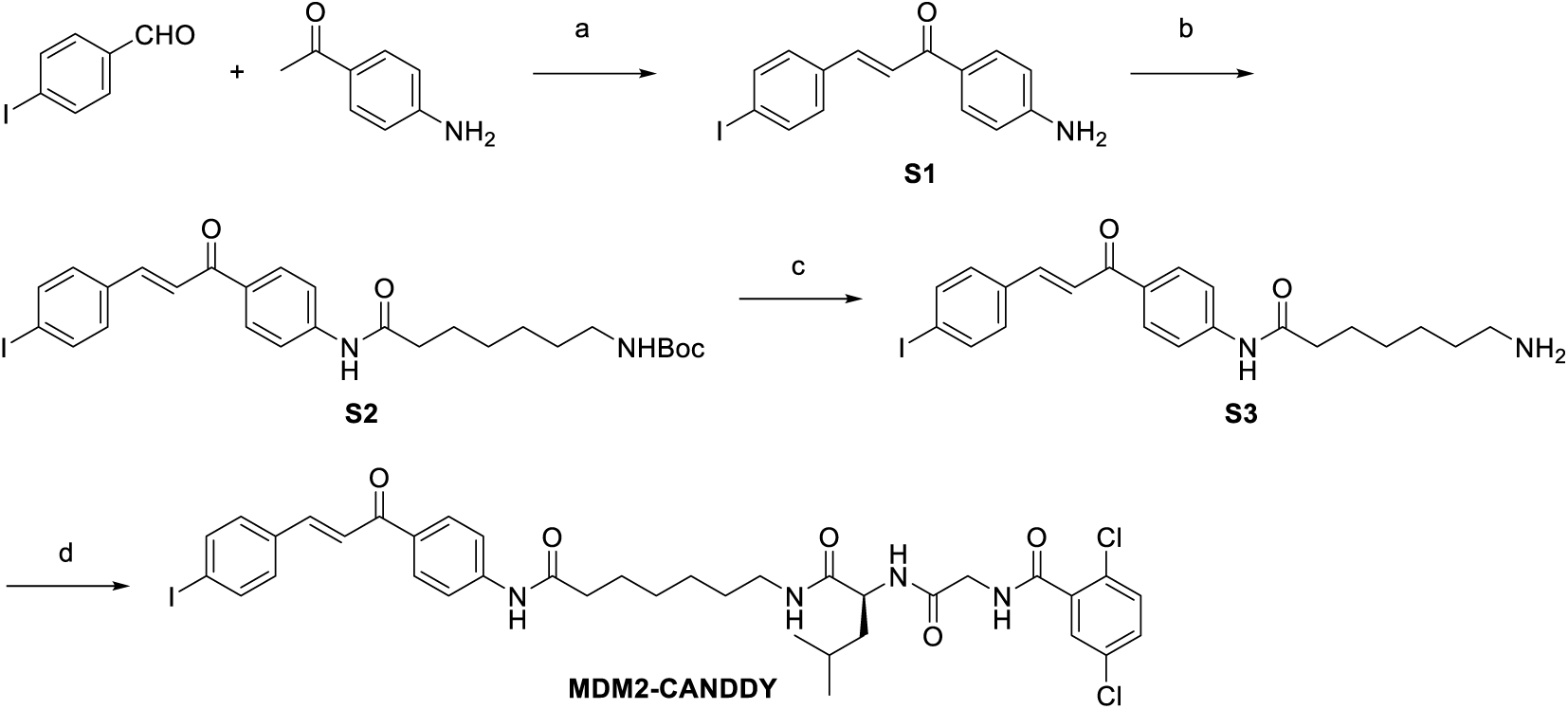
Reagent and conditions: (a) 40% KOHaq., EtOH, 0°C to rt, 60%; (b) 7-{(*tert*-Butoxycarbonyl)amino}heptanoic acid, HATU, DIPEA, DMAP, DMA∕CH_2_CI_2_, 0°C to rt, 70%; (c) TFA, CH_2_CI_2_, 0°C, 99%; (d) CANDDY-MLN, HATU, DIPEA, DMF∕CH_2_CI_2_, 0°C to rt, 65%.

To a solution of 4-Iodobenzaldehyde (7.00 g, 30.1 mmol) and 4′-Aminoacetophenone (4.08 g, 30.1 mmol) in EtOH (90 mL), 40% KOH*aq*. (60 mL) was added dropwise over 1 h at 0°C and stirred for 1 h under Ar atmosphere. After stirring for 6 h at room temperature, the resulting solution was cooled to 12°C. A precipitate was collected by filtration and washed with EtOH-water (1:2, 80 mL). The collected solid was dissolved in CHCl_3_ (800 mL) and the insoluble content was filtered and removed. The filtrate was washed with water twice, dried over Na_2_SO_4_, filtered, concentrated *in vacuo*, and thus **S1** (6.27 g, 60% yield, yellow solid) was obtained. This compound was used without further purification.

To a solution of **S1** (6.16 g, 17.7 mmol), 7-{(*tert*-Butoxycarbonyl)amino}heptanoic acid (4.35 g, 17.7 mmol), and DIPEA (4.62 mL, 26.5 mmol) in CH_2_Cl_2_-*N*,*N*-dimethyl-acetamide (DMA: 10:1, 170 mL), HATU (10.2 g, 26.8 mmol) was added at 0°C under Ar atmosphere. After stirring for 1 h at the same temperature, the solution was warmed to room temperature and stirred for 4 h, and then 4-dimethylaminopyridine (DMAP: 250 mg, 2.04 mmol) was added, and stirred further 17 h at ambient temperature. After stirring for 48 h at 30°C, the resulting solution was diluted with CH_2_Cl_2_ (1.2 L) and washed with water (0.6 L), 1N HClaq. (0.6 L) twice, sat. NaHCO_3_aq. (0.6 L) and brine (0.6 L). The combined organic layer was dried over Na_2_SO_4_, filtered, and concentrated in vacuo. The residue was purified by silica gel column chromatography (elution with 10-50% AcOEt/heptane). Next, the resulting rough solid was suspended in AcOEt (150 mL) at 50°C for 1 h and IPE (300 mL) was added, and the mixture was cooled on ice-bath. A precipitate was collected, and thus **S2** (7.14 g, 70% yield, pale yellow solid) was obtained.

To a solution of **S2** (2.50 g, 4.34 mmol) in CH_2_Cl_2_ (125 mL), TFA (63 mL) was added at 0°C under Ar atmosphere. After stirring 1.5 h at the same temperature, the resulting solution was concentrated *in vacuo*. The residue was dissolved in CH_2_Cl_2_, poured into sat. NaHCO_3_aq./water (2:1, 300 mL) and stirred for 1 h. The precipitate was collected by filtration, washed with water (20 m L) twice, dried *in vacuo*, and thus **S3** (2.05g, 99% yield, pale orange solid) was obtained.

To a solution of **CANDDY_MLN** (MLN2238 derived CANDDY tag: 0.72 g, 2.0 mmol, synthesized by WO2018092723) in DMF/CH_2_Cl_2_ (1:1, 20 mL), DIPEA (0.32 mL, 1.9 mmol) and HATU (0.76 g, 2.0 mmol) were added at 3°C. After stirring at same temperature for 30 min, **S3** (0.90g, 1.9 mmol) was added and warmed to room temperature, stirred for further 3 h at room temperature. The resulting solution was evaporated, diluted with water (200 mL), and stirred. A precipitate was collected by filtration, dried *in vacuo*, and thus **MDM2-CANDDY** (1.0 g, 65% yield, white solid) was obtained. **MDM2-CANDDY**: ^1^H-NMR (400 MHz, DMSO-d_6_) δ 0.84 (3H, d, *J* = 6.5 Hz), 0.87 (3H, d, *J* = 6.5 Hz), 1.25-1.31 (4H, m), 1.38-1.46 (4H, m), 1.53-1.59 (3H, m), 2.33 (2H, t, *J* = 7.2 Hz), 2.97-3.07 (2H, m), 3.81-3.91 (2H, m), 4.25-4.31 (1H, m), 7.52 (3H, s), 7.64 (1H, d, *J* = 15.8 Hz), 7.67 (2H, d, *J* = 8.5 Hz), 7.76 (2H, d, *J* = 8.9 Hz), 7.82 (2H, d, *J* = 8.5 Hz), 7.91-7.98 (2H, m), 7.96 (1H, d, *J* = 15.8 Hz), 8.12 (2H, d, *J* = 8.9 Hz), 8.78 (1H, dd, *J* = 5.9, 5.9 Hz), 10.23 (1H, s). HR-MS (FAB) *m*/*z*: 819.1577 (Calcd for C_37_H_42_Cl_2_IN_4_O_5_ [M+H]^+^ 819.1577).

---

**Fig. S1.**
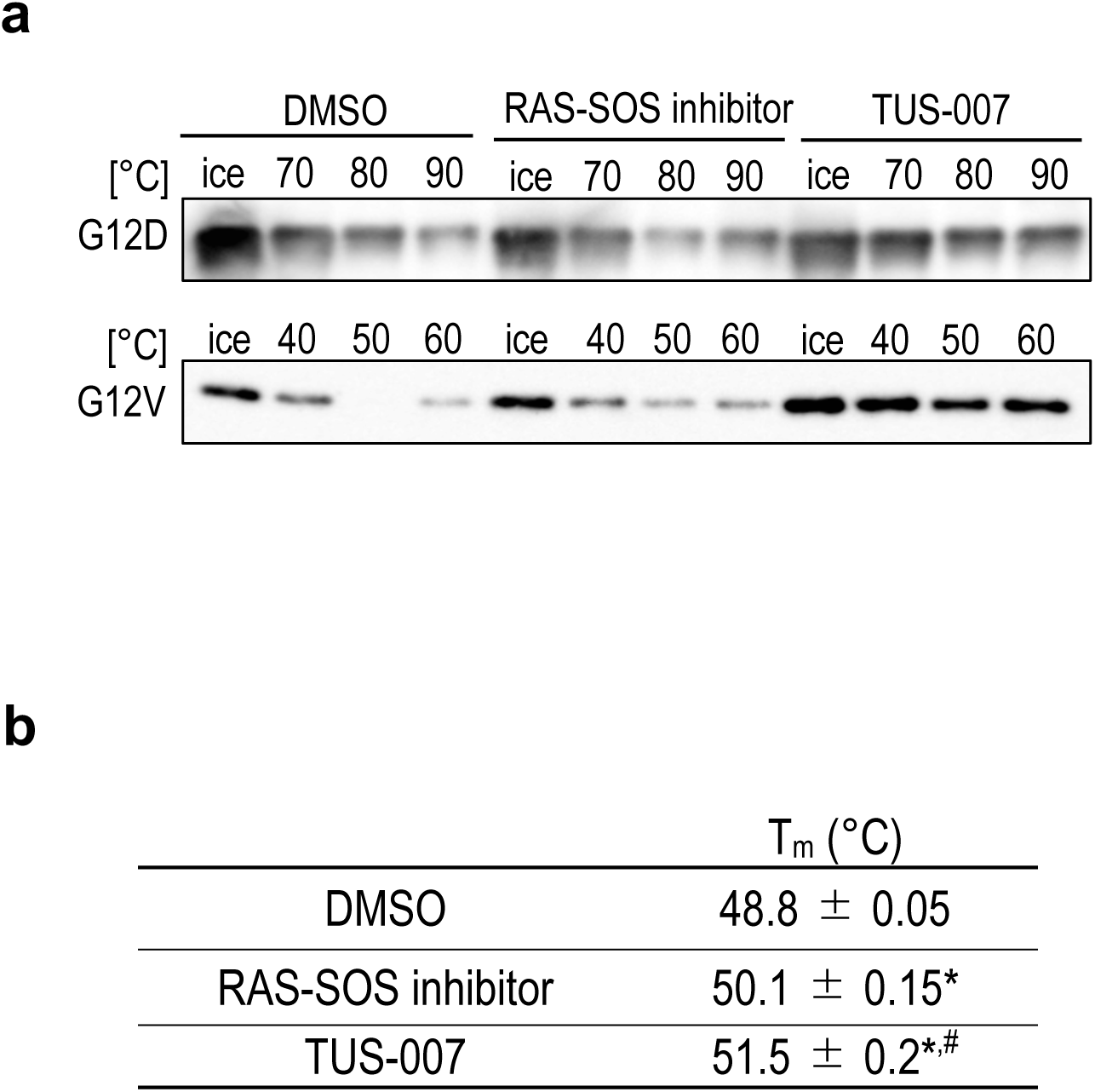
The heat resistance of KRAS G12D/V incubated with TUS-007. **a,** Evaluation of the target-binding affinity of TUS-007 to KRAS mutants by a thermal shift assay. Recombinant KRAS was treated with the vehicle, RAS-SOS inhibitor, or TUS-007 for 30 min, heated at the indicated temperature for 20 min, then analyzed by immunoblotting. **b,** The means of T_m_ values from fluorescence-based thermal shift assay shown in Fig. 1d. (mean ± SEM; n = 2). *P < 0.05 vs. DMSO, #P < 0.05 vs. RAS-SOS inhibitor.

**Fig. S2.**
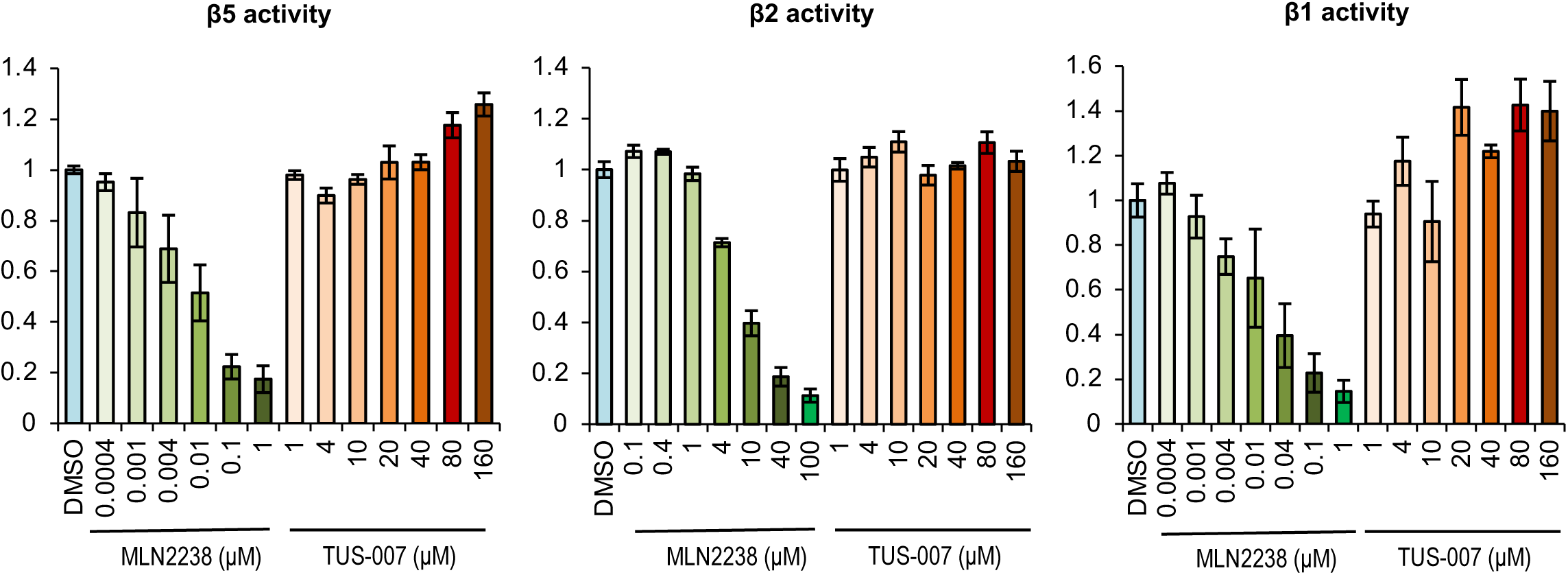
Effects of TUS-007 on proteasome activity levels. The levels of chymotrypsin-like (β5), trypsin-like (β2), and caspase-like (β1) proteasome activities was monitored by Suc-LLVY-AMC, Bz-VGR-AMC, and Z-LLE-AMC, respectively. AMC fluorescence was monitored by a plate reader with excitation and emission filters of 360 and 460 nm, respectively (DMSO, 30 min = 1) (mean ± SEM; n = 2–3).

**Fig. S3.**
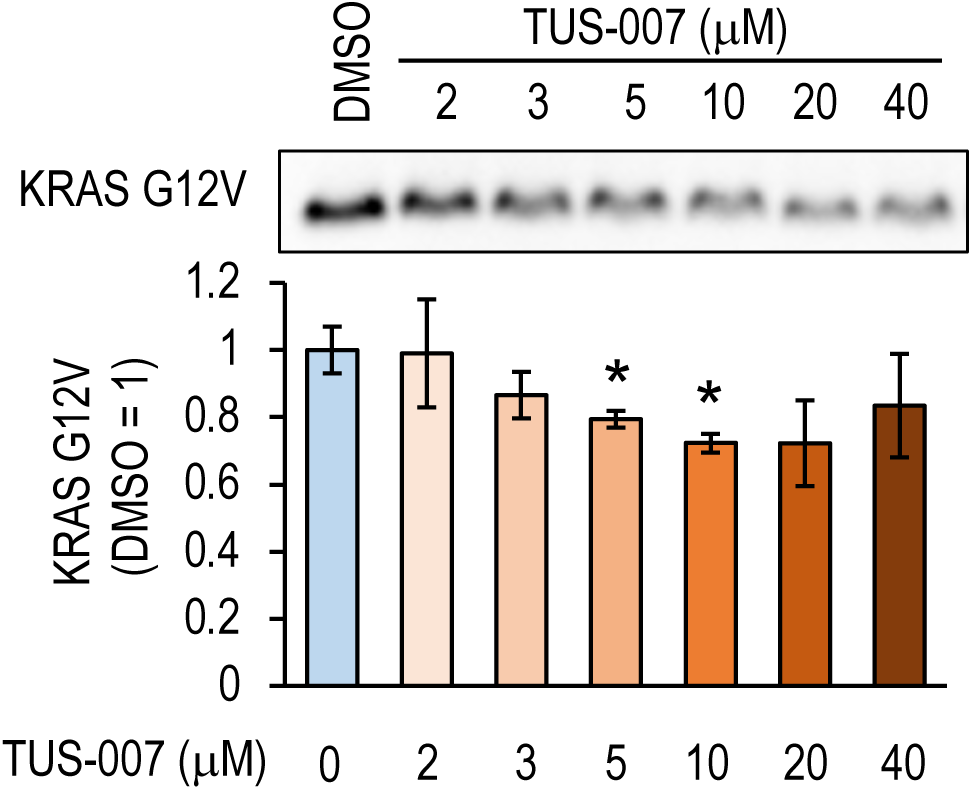
TUS-007 induced degradation of KRAS G12V in cell free assay. Successful KRAS G12V degradation by TUS-007. KRAS G12V protein was incubated with TUS-007 at the indicated concentrations in the presence of 26S proteasome for 1 h (mean ± SEM; n = 3). *P < 0.05 vs. DMSO.

**Fig. S4.**
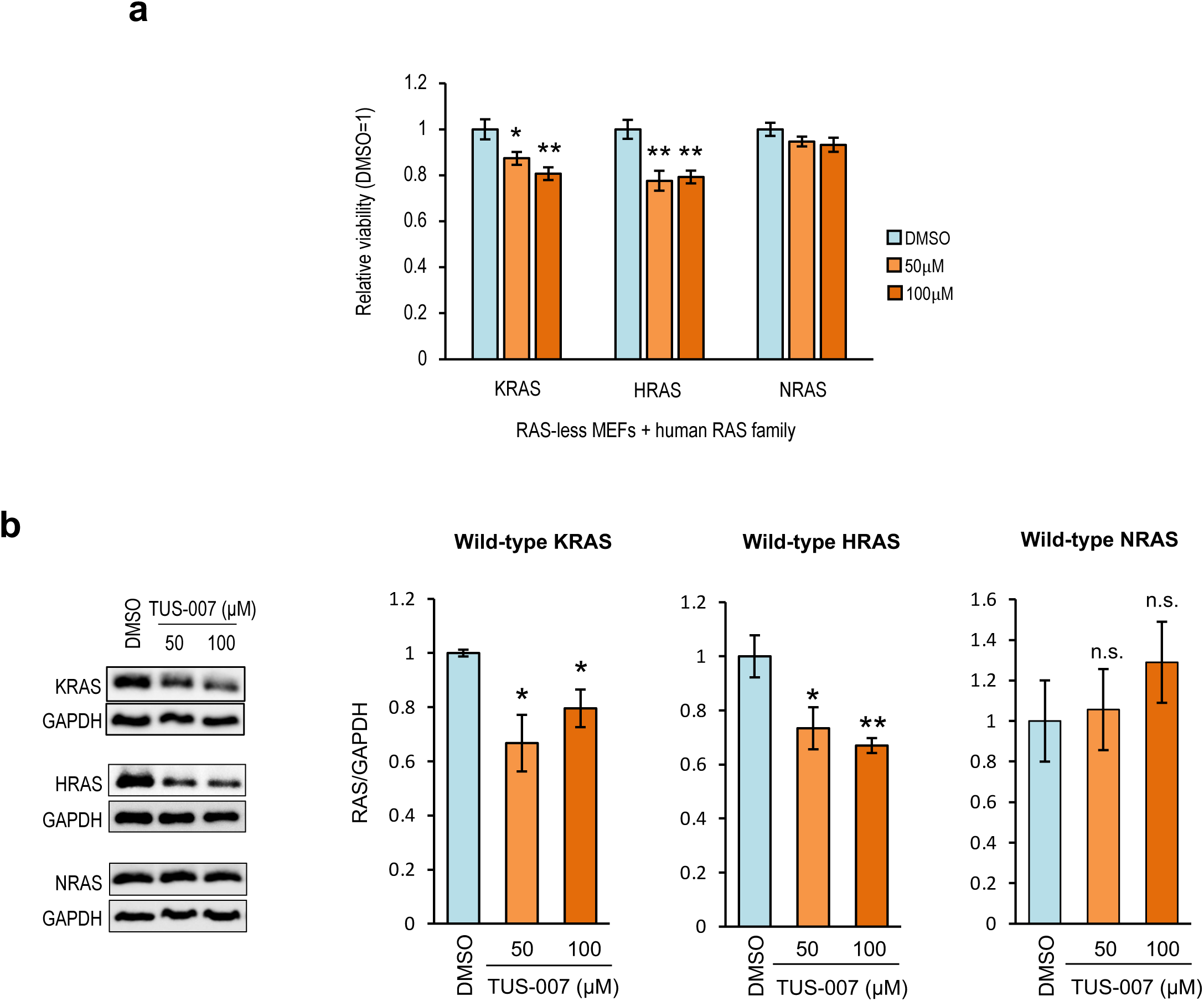
Selective chemical-knockdown of RAS variants in RAS-less MEFs expressing different types of human RAS. **a,** Relative viability of RAS-less MEFs expressing WT human RAS family members after incubation with TUS-007 or DMSO. (mean ± SEM; n = 3–5). *P < 0.05 and **P < 0.01 vs. DMSO. **b,** Degradation of WT human RAS family members in RAS-less MEFs treated with TUS-007 or DMSO for 72 h (mean ± SEM; n = 4–5). *P < 0.05 and **P < 0.01 vs. DMSO.

**Fig. S5.**
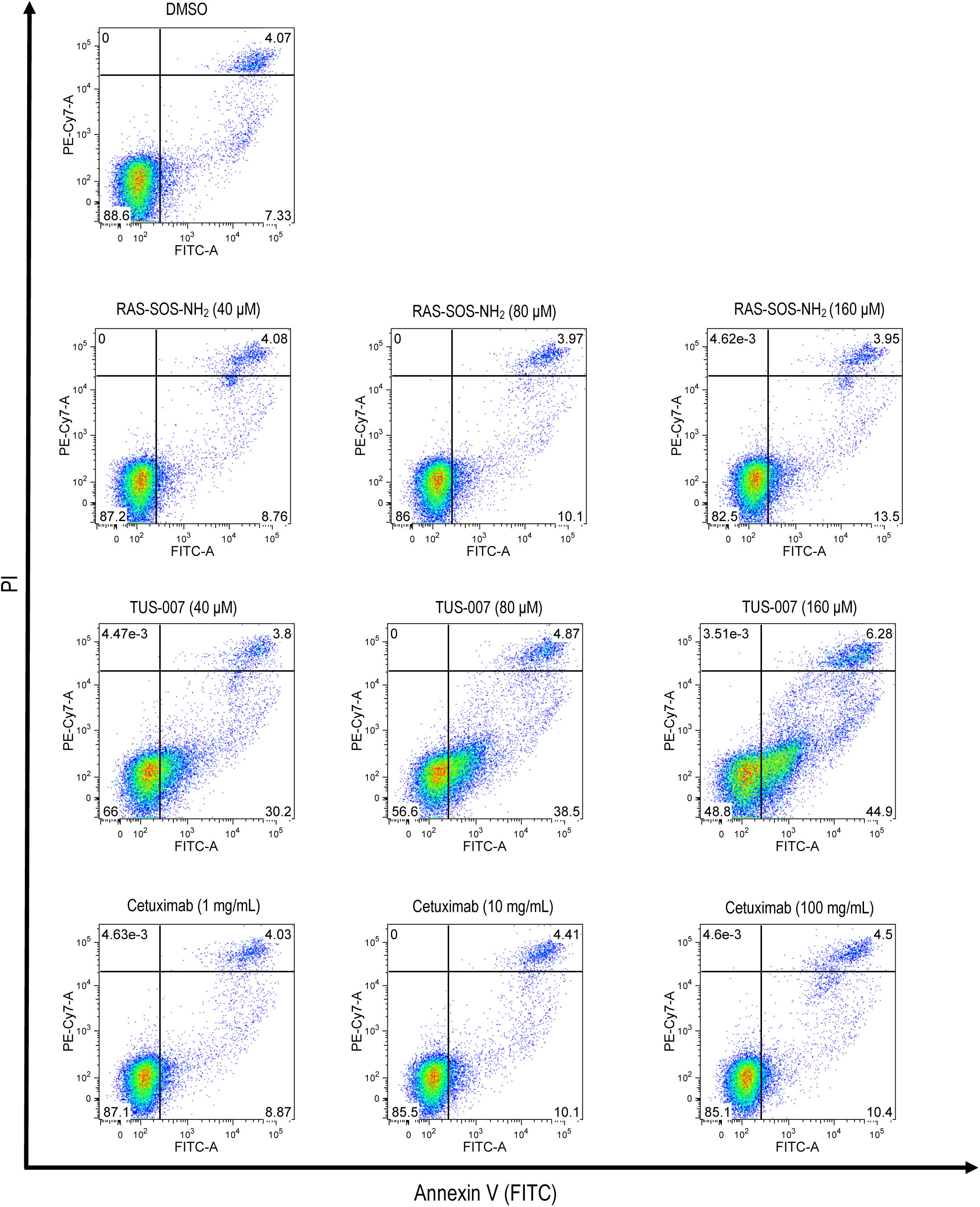
FACS plots for Annexin V-PI staining of SW620-Luc cells. SW620-Luc cells were treated with the indicated agents for 48 h, followed by staining with Annexin V-FITC and PI. The typical plots are shown.

**Fig. S6.**
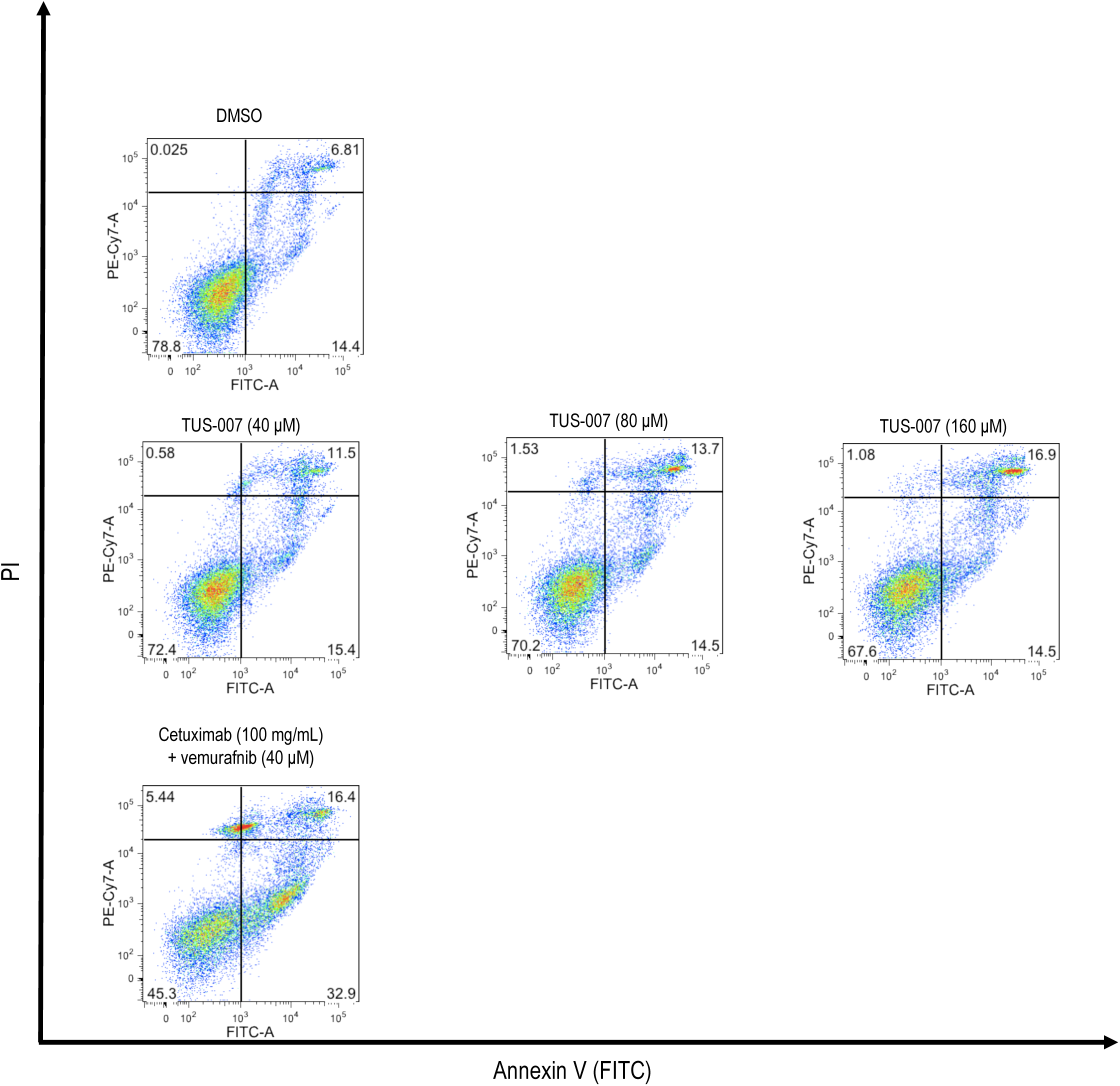
FACS plots for Annexin V-PI staining of HT29-Luc cells. HT29-Luc cells were treated with the indicated agents for 48 h, followed by staining with Annexin V-FITC and PI. The typical plots are shown.

**Fig. S7.**
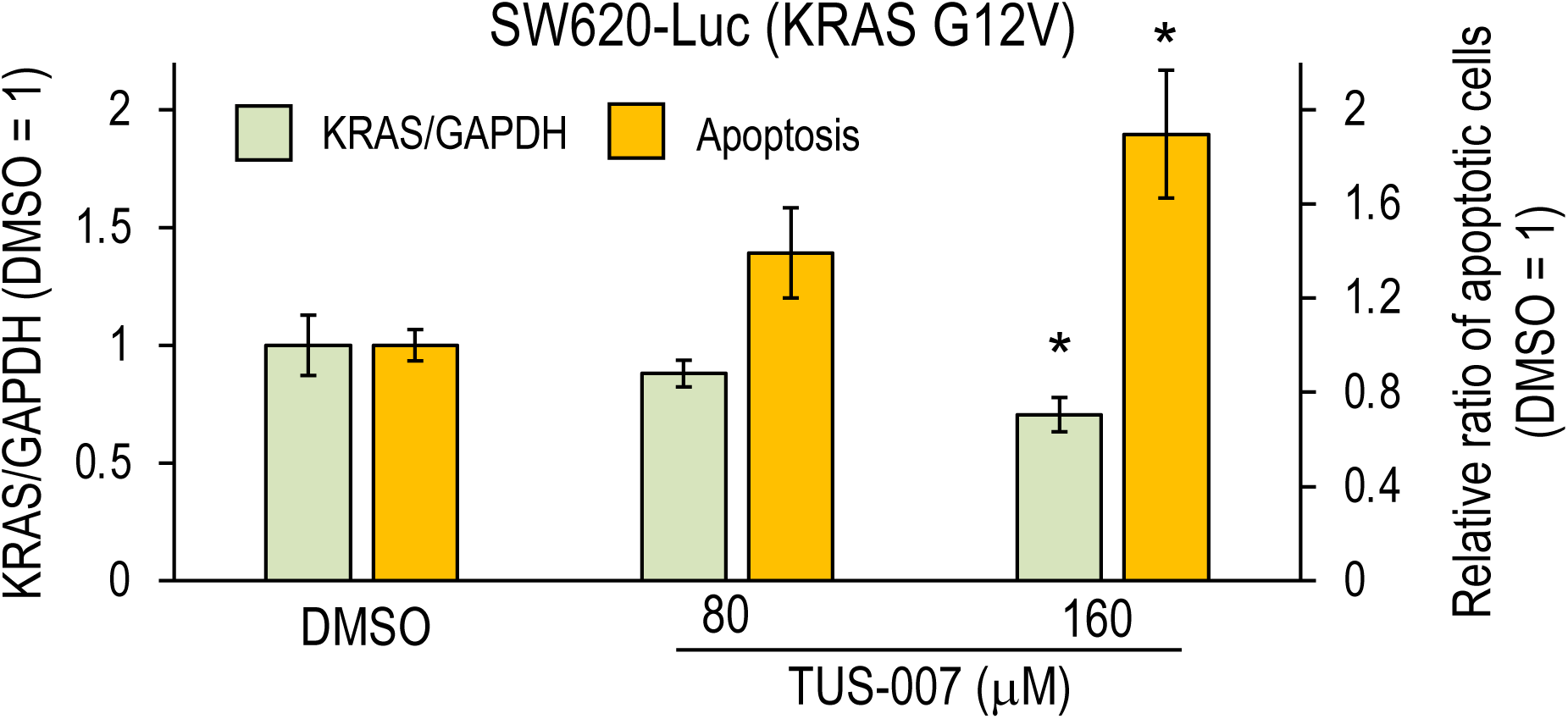
TUS-007 induced apoptosis only when KRAS degradation was detected in KRAS G12V driven cancer cells. SW620-Luc cell were treated with TUS-007 for 12 h (mean ± SEM; n = 3). The left axis indicates KRAS/GAPDH, and the right axis indicates relative ratio of apoptotic cells. *P < 0.05 vs. DMSO. The ratio of KRAS/GAPDH was determined by image analyses of immunoblotting and ration of apoptotic cells were measured using FACS.

**Fig. S8.**
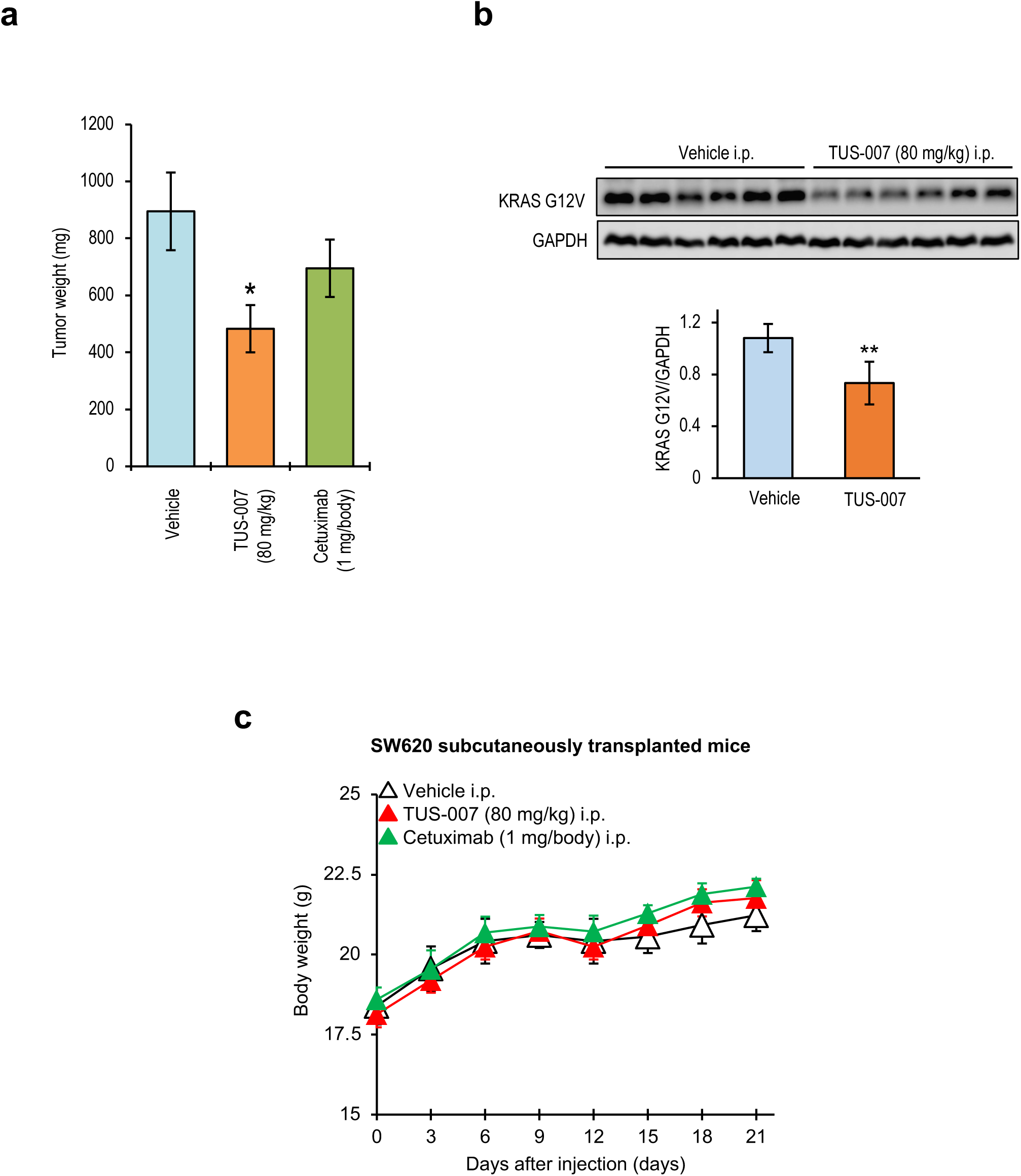
Effects of TUS-007 on the growth of colon cancer subcutaneous xenografts and toxicity. **a,** Comparison of SW620-Luc tumor weight at 21 days after injection (mean ± SEM; n = 6–8). *P < 0.05 vs. vehicle alone. **b,** Immunoblotting of KRAS G12V chemical knockdown in tumors from the same mice used in Fig. 2e (mean ± SEM; n = 6). **P < 0.01 vs. vehicle alone **c,** Body weight changes in mice with SW620-Luc cells transplanted subcutaneously (mean ± SEM; n = 6–8).

**Fig. S9.**
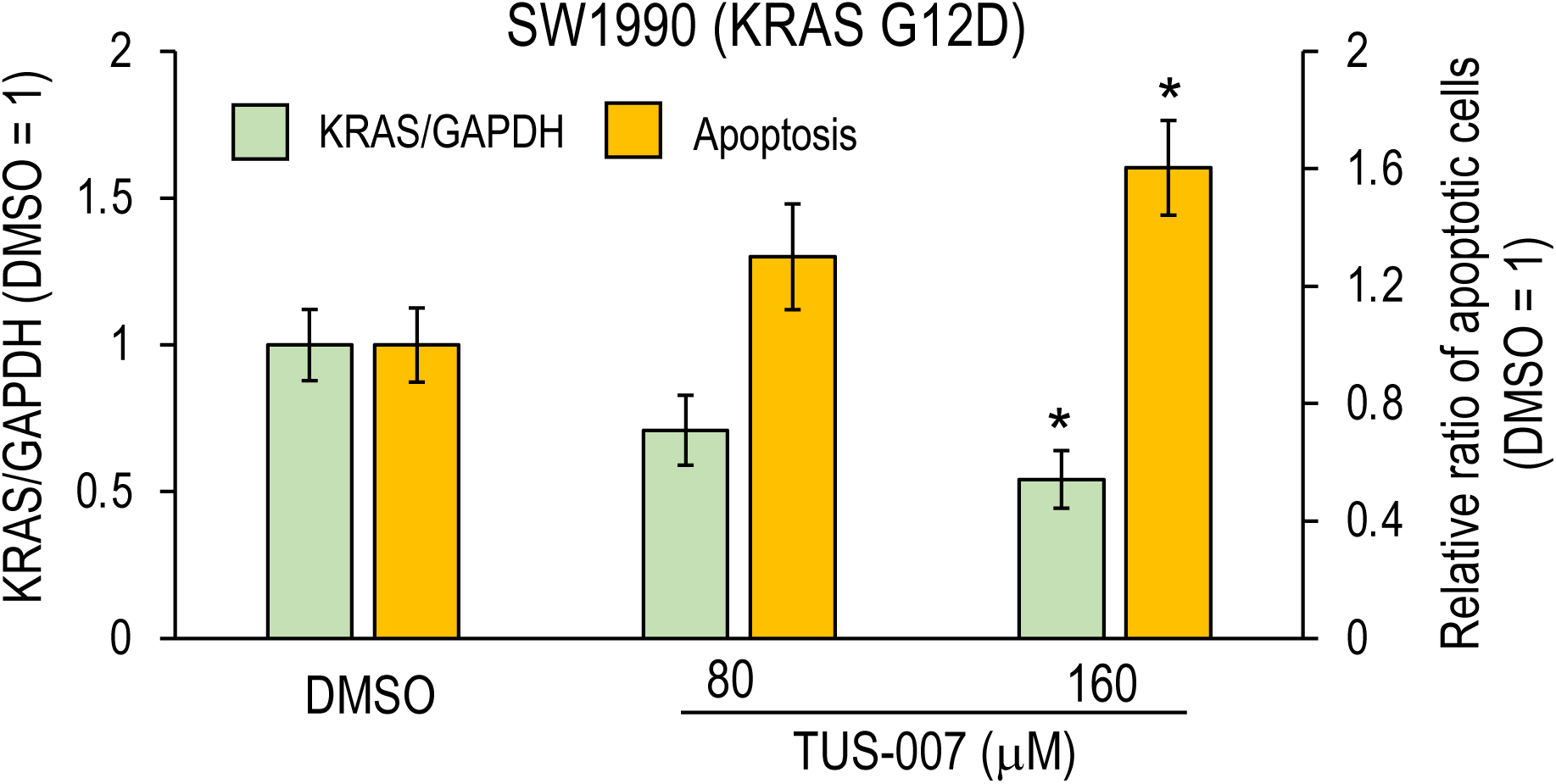
TUS-007 induced apoptosis only when KRAS degradation was detected in KRAS G12D driven cancer cells. SW1990 cell were treated with TUS-007 for 12 h (mean ± SEM; n = 3).). The left axis indicates KRAS/GAPDH, and the right axis indicates relative ratio of apoptotic cells. *P < 0.05 vs. DMSO. The ratio of KRAS/GAPDH was determined by image analyses of immunoblotting and ration of apoptotic cells were measured using FACS.

**Fig. S10.**
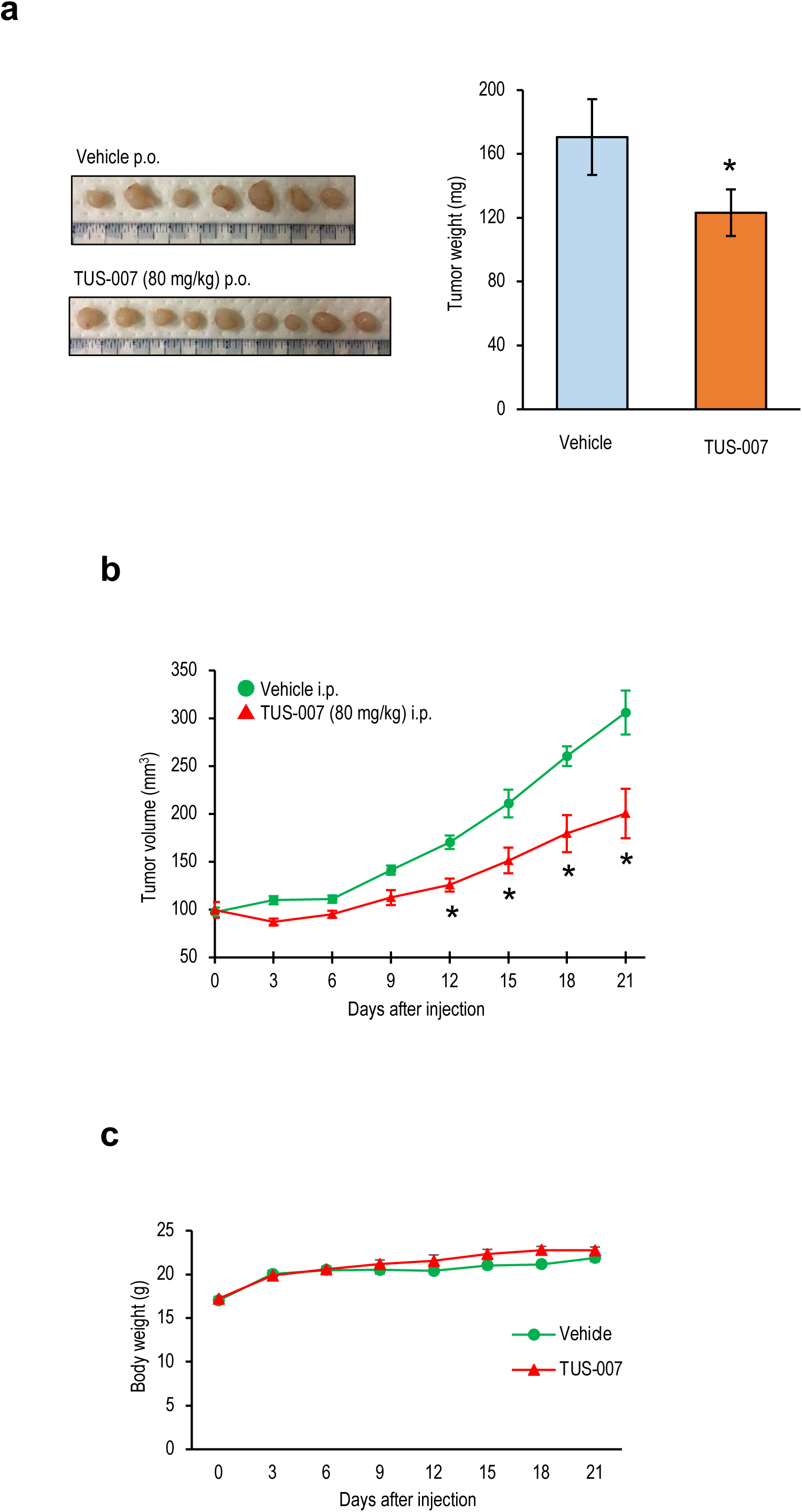
Effects of TUS-007 on the growth of pancreatic cancer subcutaneous xenografts and toxicity. **a,** Tumors of SW1990 cells from the same mice shown and comparison of tumor weight in Fig. 3e at 21 days after p.o. administration (mean ± SEM; n = 6–9). *P < 0.05 vs. vehicle alone. **b,** Tumor volume in mice with SW1990 cells transplanted subcutaneously and treated with i.p. administered TUS-007 or vehicle (mean ± SEM; n = 6–9). The agents were administered every three days. *P < 0.05 vs. vehicle. **c,** Body weight changes in mice with SW1990 cells transplanted subcutaneously (mean ± SEM; n = 6–9).

**Fig. S11.**
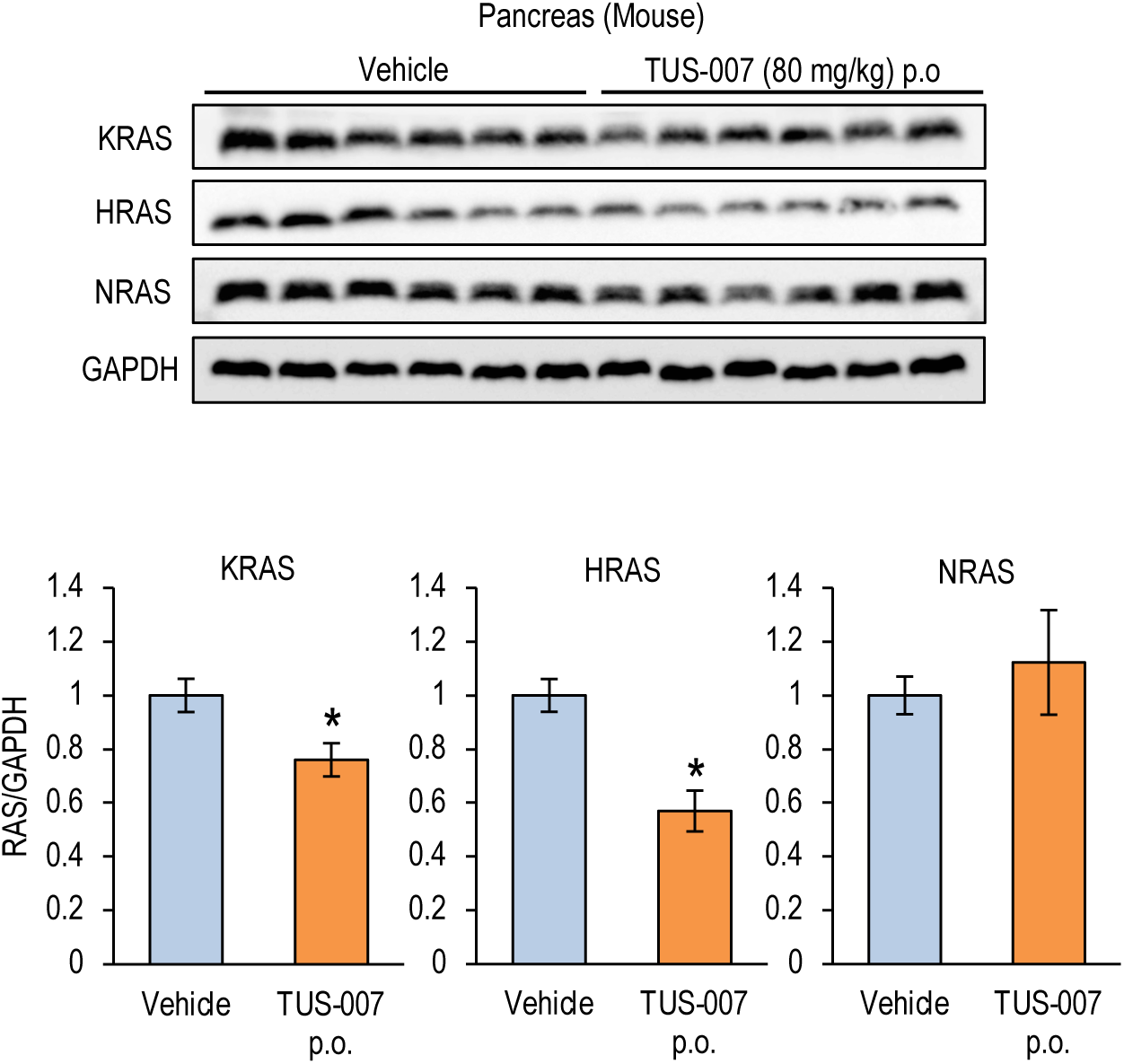
Effects of oral treatment with TUS-007 on wild type RAS in pancreas. The immunoblotting analyses of wt RAS proteins in pancreas from the same mice shown in Fig. 3e (mean ± SEM; n = 6–9). *P < 0.05 vs. vehicle alone.

**Fig. S12.**
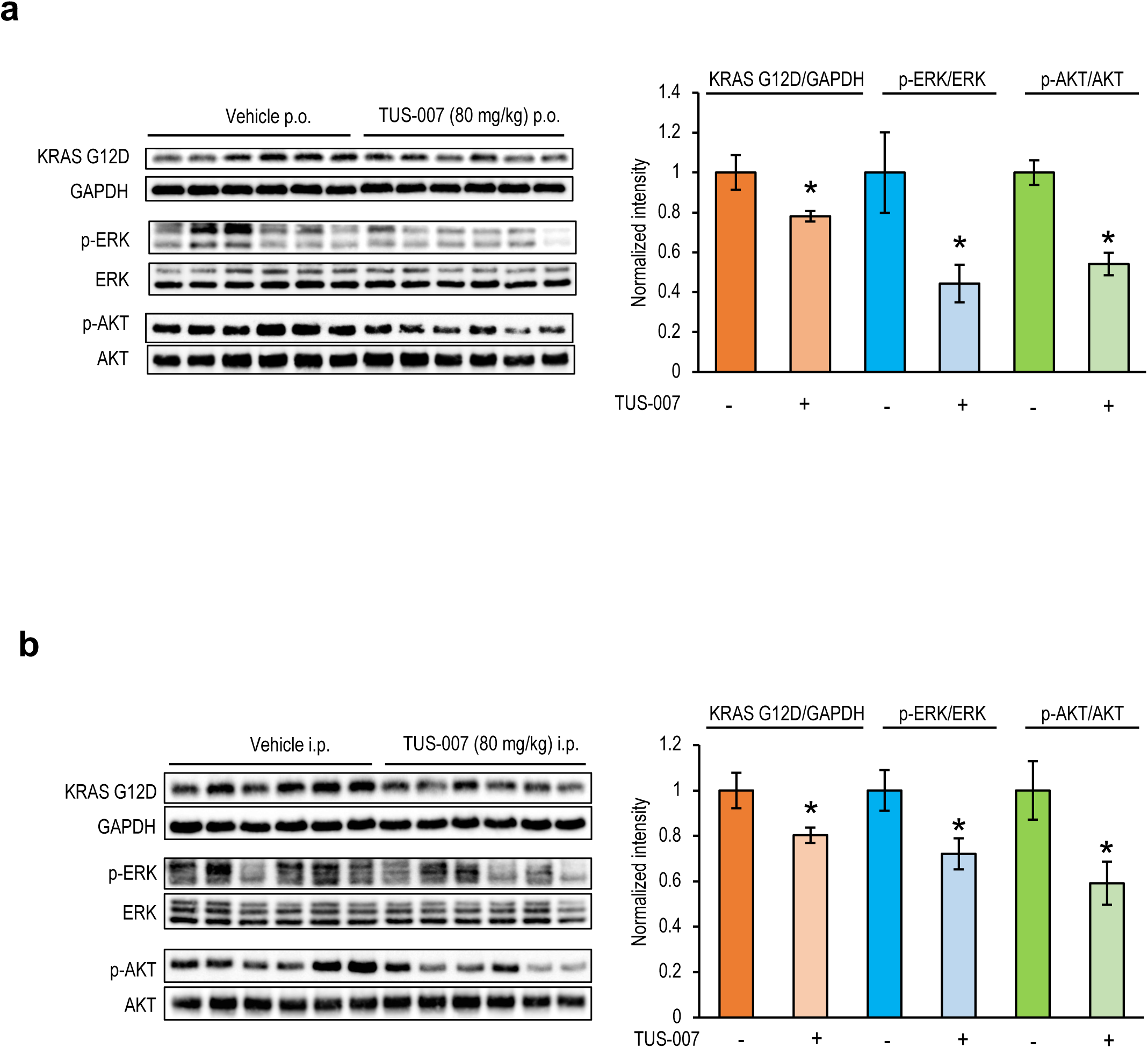
Effect of of TUS-007 on RAS signaling in pancreatic cancer subcutaneous xenografts.. Immunoblotting of KRAS and downstream signaling molecules in tumors from the same mice used in Fig 3e (**a**: p.o. treatment) and Supplementary Fig. 11c (**b**: i.p. treatment). The quantitative analysis of the immunoblotting was shown as bar graph, where the KRAS values were normalized to GAPDH, the p-ERK values were normalized to the total ERK, and the p-AKT values were normalized to the total AKT (mean ± SEM; n = 6). *P < 0.05 vs. vehicle.

**Fig. S13.**
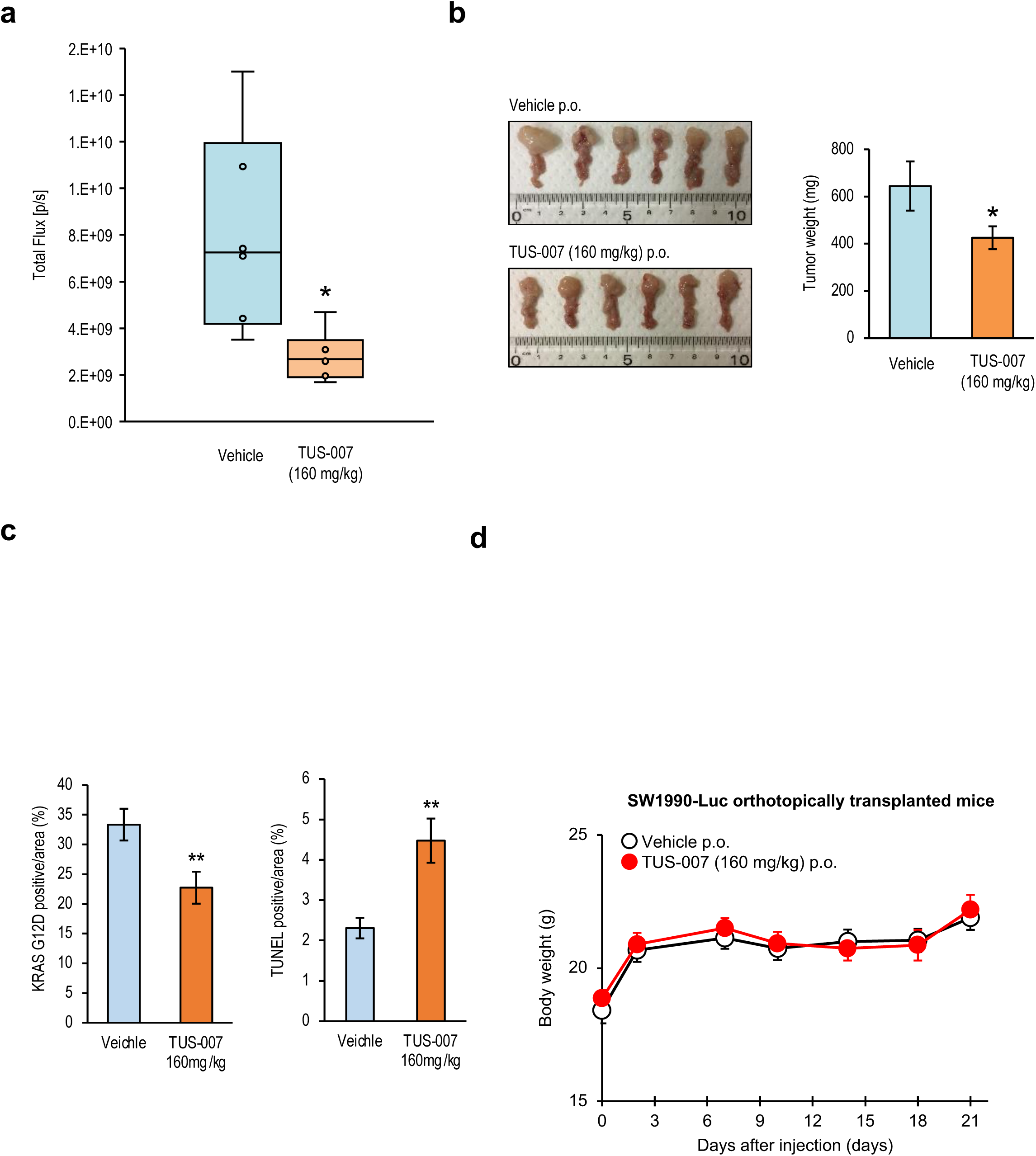
Effect of TUS-007 on growth of orthotopic pancreatic cancer xenografts expressing mutant KRAS. **a,** Tumor growth, as assessed by luciferase signal, in individual mice orthotopically transplanted SW1990-Luc cells treated with TUS-007 (red) or vehicle alone (blue) (mean ± SEM; n = 6). **b,** Pancreases from orthotopic xenograft model mice at day 21 after treatment with TUS-007 (lower-left panel) or the vehicle alone (upper-left panel) and comparison of their weights (right graph) (mean ± SEM; n = 6). *P < 0.05 vs. vehicle alone. **c,**. The quantitative analyses of the KRAS G12D-positive area (left) and TUNEL-positive area (right) in Fig. 4c (mean ± SEM; n = 6). **P < 0.01 vs. vehicle alone. **d**, Body weight changes in mice subjected to orthotopic transplantation of SW1990-Luc cells (mean ± SEM; n = 6).

**Fig. S14.**
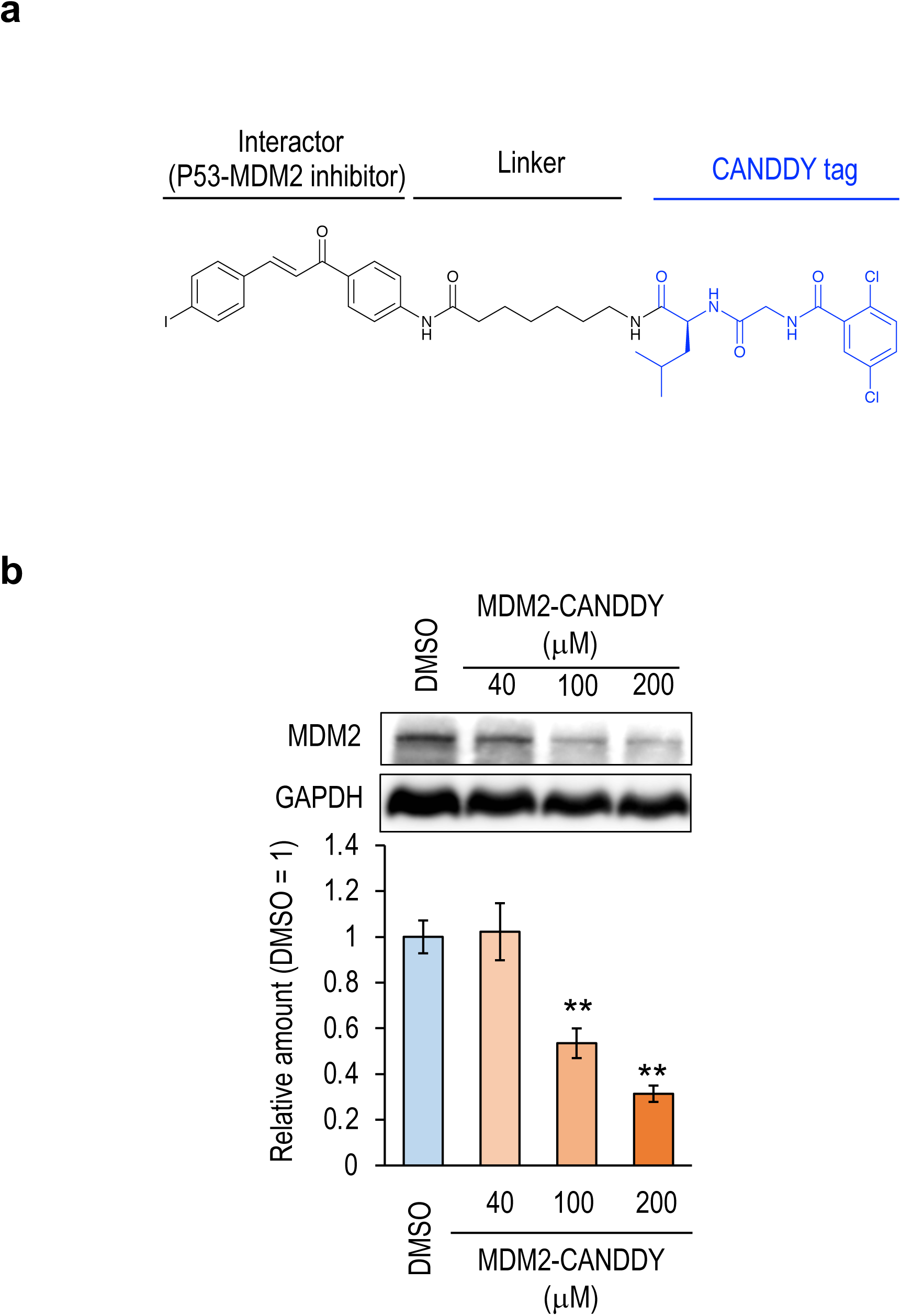
CANDDY induced degradation of MDM2, a common undruggable target. **a,** The structure of MDM2-CANDDY using P53-MDM2 PPI inhibitor, with IC50 value between 6-25 μM. **b,** MDM2-CANDDY degraded MDM2 in the dose dependent manner. The human colon cancer cells HCT-116 were incubated for 48 h with MDM2- CANDDY (mean ± SEM; n = 3). **P < 0.01 vs. DMSO.

**Table S1.** The concentration of TUS-007 in pancreas was maintained for 72 h. The concentration of TUS-007 in pancreas from healthy wild-type mice i.p. treated with TUS-007 at 80 mg/kg (means ± SEM; n = 5). The samples were obtained at indicated time points after i.p. injection. The concentration of TUS-007 was measured with HPLC.

| Hours after i.p | 1h | 20h | 72h |
| --- | --- | --- | --- |
| TUS-007 in pancreas<br>(ng/mg) | 45.7 ± 20.3 | 55.0 ± 28.6 | 61.8 ± 30.0 |

**Table S2.**
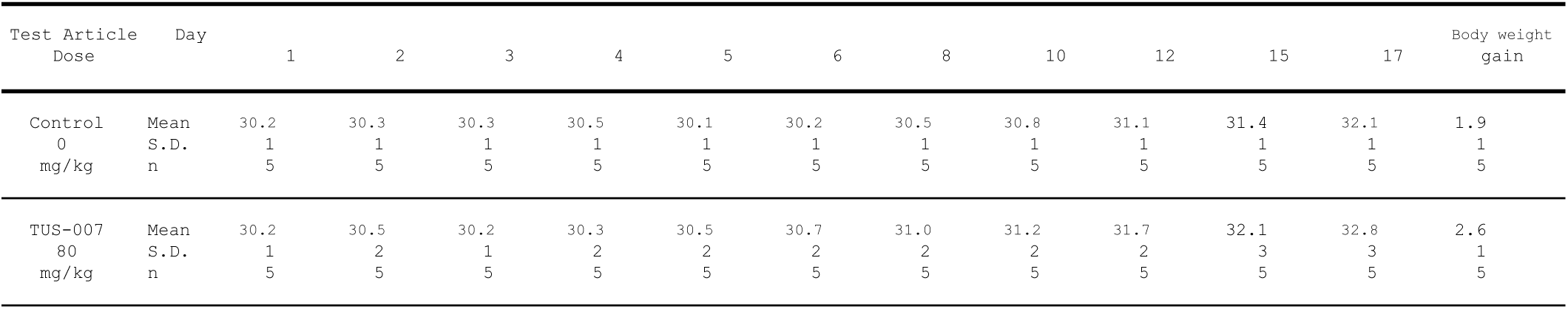
Body weight change in toxicological tests. TUS-007 did not affected on the body weight changes of mice.

**Table S3.** Weight of organs in toxicological tests. TUS-007 did not affected on the organ weight of mice.

| Test Article<br>Dose |  | F.B.W.@@<br>g | Heart<br>mg | Lung<br>mg | Liver<br>g | Spleen<br>mg | Kidney-RL<br>mg |
| --- | --- | --- | --- | --- | --- | --- | --- |
| Control<br>0<br>mg/kg | Mean | 32.5 | 147 | 175 | 1.81 | 96 | 448 |
|  | S.D. | 0.9 | 9 | 10 | 0.14 | 17 | 34 |
|  | n | 5 | 5 | 5 | 5 | 5 | 5 |
| TUS-007<br>80<br>mg/kg | Mean | 33.0 | 157 | 184 | 1.78 | 85 | 483 |
|  | S.D. | 2.6 | 15 | 21 | 0.22 | 22 | 68 |
|  | n | 5 | 5 | 5 | 5 | 5 | 5 |

| Test Article<br>Dose |  | Heart<br>mg/100g | Lung<br>mg/100g | Liver<br>g/100g | Spleen<br>mg/100g | Kidney-RL<br>mg/100g |
| --- | --- | --- | --- | --- | --- | --- |
| Control<br>0<br>mg/kg | Mean | 451 | 538 | 5.57 | 295 | 1378 |
|  | S.D. | 26 | 19 | 0.37 | 45 | 87 |
|  | n | 5 | 5 | 5 | 5 | 5 |
| TUS-007<br>80<br>mg/kg | Mean | 475 | 557 | 5.38 | 255 | 1462 |
|  | S.D. | 16 | 37 | 0.28 | 52 | 123 |
|  | n | 5 | 5 | 5 | 5 | 5 |

**Table S4.** Histological observations of organs in toxicological tests. TUS-007 did not cause histological changes in mice.

| Organs | Sex: | M | M |
| --- | --- | --- | --- |
|  | Group: | Control | TSU-007 |
|  | Dose (mg/kg) : | 0 | 80 |
| Findings | Number: | 5 | 5 |
| Brain(cerebrum) |  |  |  |
| Number examined |  | 5 | 5 |
| Not remarkable |  | 5 | 5 |
| Heart |  |  |  |
| Number examined |  | 5 | 5 |
| Not remarkable |  | 5 | 5 |
| Necrosis/infiltrate |  | 0 | 0 |
| minimal |  | 0 | 0 |
| Intestine,rectum |  |  |  |
| Number examined |  | 5 | 5 |
| Not remarkable |  | 5 | 5 |
| Kidney |  |  |  |
| Number examined |  | 5 | 5 |
| Not remarkable |  | 5 | 5 |
| Cyst |  | 0 | 0 |
| minimal |  | 0 | 0 |
| Liver |  |  |  |
| Number examined |  | 5 | 5 |
| Not remarkable |  | 5 | 5 |
| Infiltrate,neutrophil |  | 0 | 0 |
| minimal |  | 0 | 0 |
| Necrosis,single cell,hepatocyte |  | 0 | 0 |
| minimal |  | 0 | 0 |
| Lung (bronchus) |  |  |  |
| Number examined |  | 5 | 5 |
| Not remarkable |  | 5 | 5 |
| Pancreas |  |  |  |
| Number examined |  | 5 | 5 |
| Not remarkable |  | 5 | 5 |
| Spleen |  |  |  |
| Number examined |  | 5 | 5 |
| Not remarkable |  | 5 | 5 |

**Table S5.** Hematology in toxicological tests. TUS-007 did not cause hematological changes in mice.

| Test Article<br>Dose | | RBC<br>10E4/μL | HGB<br>g/dL | HCT<br>% | MCV<br>fL | MCH<br>pg | MCHC<br>g/dL | Retic<br>10E9/L | PLT<br>10E4/μL |
| --- | --- | --- | --- | --- | --- | --- | --- | --- | --- |
| Control<br>0<br>mg/kg | Mean | 881 | 13.5 | 47.2 | 53.5 | 15.3 | 28.7 | 241.4 | 113.2 |
|  | S.D. | 59 | 0.7 | 2.9 | 0.9 | 0.5 | 0.8 | 26.9 | 7.0 |
|  | n | 5 | 5 | 5 | 5 | 5 | 5 | 5 | 5 |
| TUS-007<br>80<br>mg/kg | Mean | 876 | 13.3 | 46.4 | 53.1 | 15.2 | 28.6 | 281.2 | 120.8 |
|  | S.D. | 53 | 0.4 | 1.4 | 2.7 | 0.8 | 0.4 | 70.7 | 3.0 |
|  | n | 5 | 5 | 5 | 5 | 5 | 5 | 5 | 5 |

| Test Article<br>Dose | | WBC<br>10E2/μL | LYMP<br>10E2/μL | NEUT<br>10E2/μL | EOS<br>10E2/μL | BASO<br>10E2/μL | MONO<br>10E2/μL | LUC<br>10E2/μL |
| --- | --- | --- | --- | --- | --- | --- | --- | --- |
| Control<br>0<br>mg/kg | Mean | 42.5 | 32.3 | 7.8 | 1.4 | 0.0 | 0.9 | 0.0 |
|  | S.D. | 6.2 | 3.9 | 3.2 | 0.6 | 0.1 | 0.3 | 0.1 |
|  | n | 5 | 5 | 5 | 5 | 5 | 5 | 5 |
| TUS-007<br>80<br>mg/kg | Mean | 37.5 | 28.3 | 6.8 | 1.1 | 0.0 | 1.2 | 0.1 |
|  | S.D. | 14.2 | 12.0 | 3.0 | 0.4 | 0.1 | 0.7 | 0.1 |
|  | n | 5 | 5 | 5 | 5 | 5 | 5 | 5 |

**Table S6.**
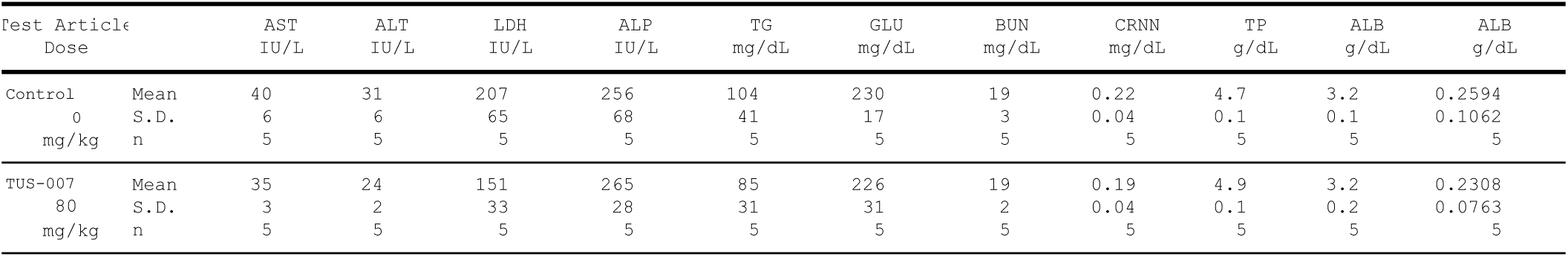
Biochemical examinations in toxicological tests. TUS-007 did not cause biochemical changes in mice.

